# Paradoxical Phenotype of Fibromyalgia Neutrophils with Elevated Baseline Inflammation but Blunted Response to Stimulation

**DOI:** 10.1101/2025.09.15.676281

**Authors:** Sahel Jahangiri Esfahani, Anahita Oveisi, Lucas Vasconcelos Lima, Sara Caxaria, Marc Parisien, Julia Vignone, Francesca Montagna, Shafaq Sikandar, Carolina Beraldo Meloto, Jeffrey S. Mogil, Luda Diatchenko

## Abstract

Fibromyalgia (FM) is a severe pain condition of unknown etiology. Here, we performed transcriptomics analyses of peripheral neutrophils exposed to an inflammatory stimulus, comparing responses of neutrophils obtained from FM patients versus healthy controls. We observed a state of inflammation in neutrophils from FM patients. However, FM neutrophils were unable to efficiently respond to lipopolysaccharide (LPS). This impairment was especially characteristic of FM patients with no improvement after 5 years after diagnosis in comparison with those who did improve. Blood plasma from FM patients directly stimulated a wide range of primary sensory neurons *in vitro* and induced pain hypersensitivity when injected into mice. Further analysis identified NF-κB suppression as a key biological process associated with low-grade inflammation and LPS non-responsiveness in neutrophils from FM patients. The clinically used NF-κB activator, bryostatin, alleviated hypersensitivity in mice treated with FM plasma, pointing to controlled inflammation induction through reactivation of the NF-κB pathway as a possible therapeutic target for FM treatment. Our whole blood single-cell RNA sequencing replicated this NF-κB-driven inflammation observed in bulk analyses transcriptomics in FM patients and revealed that this inflammatory signature is strongly pronounced not only in neutrophils, but across a broad range of immune cells.

## INTRODUCTION

Chronic pain adversely affects approximately 20% of the global population and imposes a significant economic burden exceeding $600 billion annually just in the United States (*1*). Fibromyalgia (FM), an idiopathic chronic pain condition, is characterized by widespread musculoskeletal pain, fatigue, negative mood, and cognitive dysfunction, affecting 2–8% of the population, with women disproportionately affected at a 9:1 ratio (*2, 3*). Current treatments for FM encompass multidisciplinary approaches, combining pharmacological interventions, such as analgesics and antidepressants (*4*), with non-pharmacological strategies, including cognitive-behavioural therapy and exercise (5–7). However, the effectiveness of these non-specific treatments varies, and achieving long-term symptom relief remains a considerable challenge. Despite relatively high disease prevalence and profound health burdens imposed by FM, its molecular underpinnings remain elusive, necessitating a comprehensive exploration to enhance diagnostic accuracy and therapeutic efficacy (*4*).

FM is increasingly recognized as a heterogeneous condition with no cure, with inconsistent reports on the rate of symptom improvement over time (*8, 9*). The few existing studies indicate a 20–47% improvement rate over 1–2 years among people with FM (8–10), although the mechanisms driving this improvement are unknown. Increasing evidence suggests the importance of the immune system in chronic pain development and resolution (11–14). Cytokine profile and immune system abnormalities have been reported in the peripheral blood of people with FM in multiple studies (*15*). One of these features, a low-grade inflammation profile, has been repeatedly reported in a range of chronic pain conditions, including FM (*16, 17*). Furthermore, recent research has highlighted the pivotal causal role of FM neutrophils in the pathogenesis of the disease (*18*). On the other hand, acute inflammatory response driven by neutrophils has been recently shown as a *protective* immune mechanism, preventing chronic pain development in acute pain patients (*19*). These apparently contradictory observations underscore the imperative for a more comprehensive and detailed examination of these immune cells (*18*).

Here, we undertook multidimensional transcriptomics analyses on peripheral neutrophils obtained from patients with FM and healthy controls to evaluate their response to the immune stimulator, lipopolysaccharide (LPS). We then validated our findings in animal models, demonstrating the pivotal role of inflammation suppression through inhibition of the nuclear factor kappa-light-chain-enhancer of activated B cells (NF-κB) transcription factor (TF) in disease persistence, and its reactivation as a new potential therapeutic target for FM treatment. We further validated our results in whole blood single-cell RNA sequencing (scRNA-seq) of the independent FM cohort.

## RESULTS

### STUDY DESIGN AND COHORT DESCRIPTION

Blood samples and clinical data were collected from 64 healthy controls (HC) and 62 patients with fibromyalgia (FM). Neutrophils were isolated from whole blood and cultured under two conditions, unstimulated and LPS-stimulated. Following RNA extraction, bulk RNA sequencing was performed on all samples (Figure. 1A). The demographic and clinical characteristics of a total of 126 participants in the study cohort are summarized in Table 1. There were no significant group differences in age, sex, or BMI. In contrast, FM patients scored significantly higher on all symptom and psychological measures including the Fibromyalgia Survey, Beck Anxiety and Depression Inventories, Fibromyalgia Impact Questionnaire, Symptom Checklist-90-R, Pittsburgh Sleep Quality Index and Perceived Stress Scale, and they also exhibited lower pressure pain thresholds compared to controls (all p-values < 0.05).

**Figure. 1.**
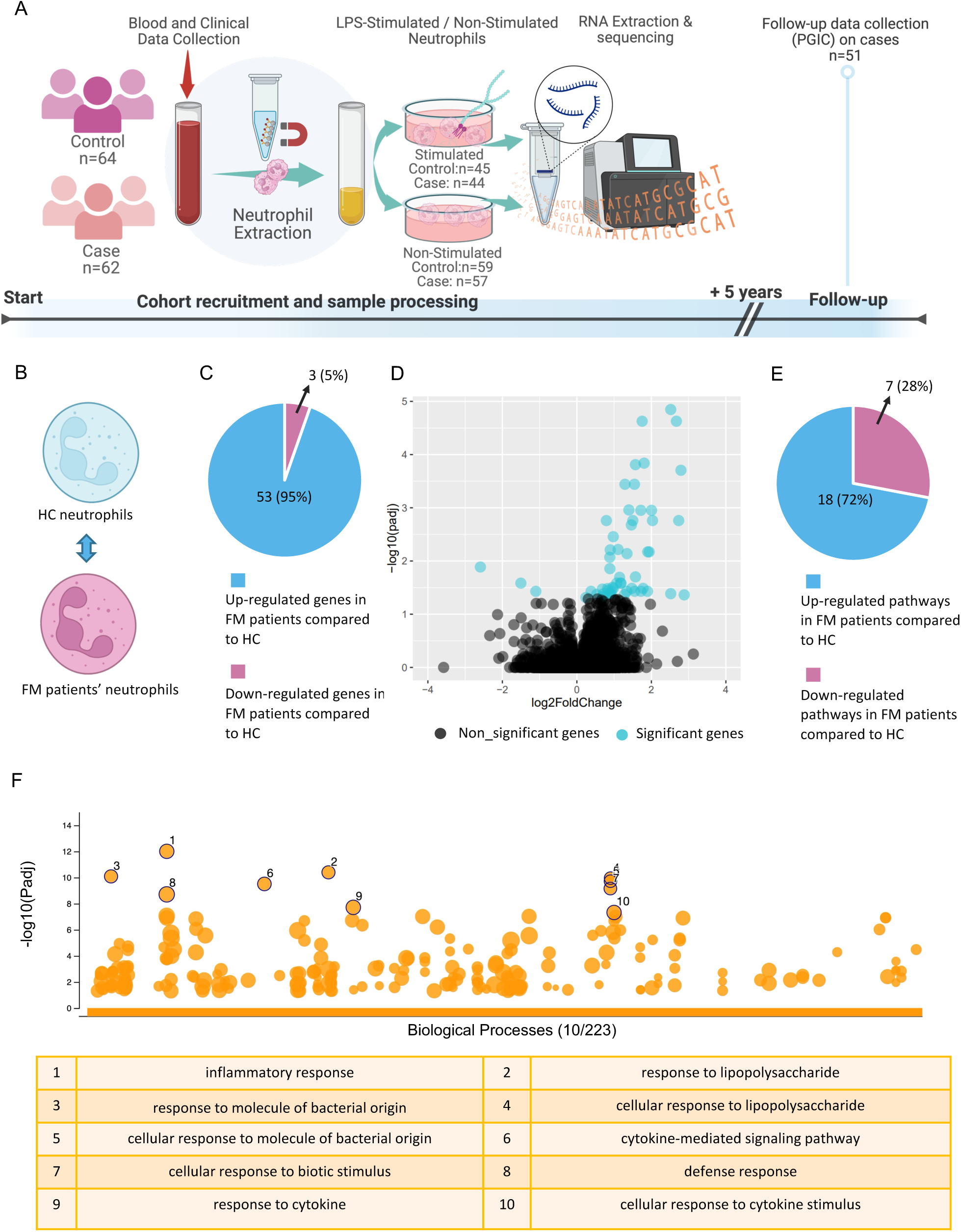
Neutrophils from FM patients show activated phenotype. (A) Schematic representing the human cohort study design and follow-up. (B) Schematic representing the design of the experimental groups in the analysis. (C) Pie chart representing the counts for the number of differentially expressed genes (DEG) in neutrophils from FM patients versus HC. (D) Scatterplot representing the log2fold changes versus the statistical significance of the DEGs in neutrophils from FM patients versus HC. Black symbols represent genes regulated with an adjusted p-value of higher than 0.05, and blue symbols lower than p-value=0.05. (E) Pie chart representing the statistics of Fgsea pathway analysis on DEGs in neutrophils from FM patients versus HC. (F) Bubble chart representing gProfiler pathway analysis results performed on the significant DEGs in neutrophils from FM patients and HC. The x-axis represents the biological processes from GO. The y-axis shows the adjusted enrichment p-values in negative log10 scale.

**Table 1.** Demographic and clinical characteristics of the fibromyalgia patients and controls. N-Miss represents the number of missing values per group. p-values represent group comparisons (Mann–Whitney U test for continuous variables; Fisher’s exact test for categorical variables). Asterisks (*) denote statistically significant results.

|  | Controls (n = 64) | Cases (n = 62) | Total (N = 126) | p-value |
| --- | --- | --- | --- | --- |
| **Age** |  |  |  | 0.059 |
| Mean (SD) | 53.2 (9.2) | 55.2 (7.1) | 54.2 (8.27) |  |
| Median (range) | 51.5 (40.0 - 77.0) | 55.5 (40.0 - 73.0) | 54.5 (40.0 - 77.0) |  |
| **Sex** |  |  |  | 0.076 |
| Female | 54 (84.4%) | 59 (95.2%) | 113 (89.7%) |  |
| Male | 10 (15.6%) | 3 (4.8%) | 13 (10.3%) |  |
| **BMI** |  |  |  | 0.120 |
| N-Miss | 0 | 0 | 0 |  |
| Mean (SD) | 26.8 (5.5) | 28.9 (7.2) | 27.8 (6.5) |  |
| Median (range) | 26.6 (17.9 - 40.9) | 27.8 (18.8 - 57.1) | 27.45 (17.9 - 57.1) |  |
| **Fibromyalgia Survey (FMS)** |  |  |  | 3.17E-21* |
| N-Miss | 0 | 1 | 1 |  |
| Mean (SD) | 4.9 (4.4) | 21.7 (5) | 13.1 (9.7) |  |
| Median (range) | 3.5 (0 - 19) | 22 (11 - 30) | 12 (0 - 30) |  |
| **Beck Anxiety Inventory** |  |  |  | 2.50E-16* |
| N-Miss | 1 | 0 | 1 |  |
| Mean (SD) | 4 (4.4) | 18.1 (8.9) | 10.9 (9.9) |  |
| Median (range) | 3 (0 - 18) | 18 (1 - 40) | 7 (0 - 40) |  |
| **Fibromyalgia Impact Questionnaire** |  |  |  | 4.35E-23* |
| N-Miss | 0 | 1 | 1 |  |
| Mean (SD) | 6.2 (8.5) | 57.8 (18.6) | 30.1 (29.4) |  |
| Median (range) | 2.9 (0 - 42.7) | 54.9 (22.4 - 93.3) | 22.1 (0 - 93.3) |  |
| **Symptom Checklist 90 R** |  |  |  | 2.80E-21* |
| N-Miss | 1 | 0 | 1 |  |
| Mean (SD) | 0.6 (0.6) | 3.9 (1.2) | 2.2 (1.9) |  |
| Median (range) | 0.4 (0 - 2.14) | 4 (1.1 - 6.3) | 1.7 (0 - 6.3) |  |
| **Pressure Pain Threshold Score** |  |  |  | 2.23E-05* |
| N-Miss | 2 | 4 | 6 |  |
| Mean (SD) | 4.6 (1.9) | 4.5 (7.0) | 4.6 (5.0) |  |
| Median (range) | 4.8 (1.3 - 10.8) | 2.7 (0.3 - 34.1) | 3.7 (0.3 - 34.1) |  |
| **Pittsburgh Sleep Quality Index score** |  |  |  | 1.18E-12* |
| N-Miss | 1 | 0 | 1 |  |
| Mean (SD) | 5 (3.1) | 10.7 (4.2) | 7.8 (4.7) |  |
| Median (range) | 5 (1 - 16) | 10 (2 - 20) | 7 (1 - 20) |  |
| **Perceived Stress Scale** |  |  |  | 0.03* |
| N-Miss | 1 | 0 | 1 |  |
| Mean (SD) | 30.5 (6.4) | 33.4 (4) | 31.9 (5.5) |  |
| Median (range) | 32 (0 - 40) | 33 (27 - 43) | 33 (0 - 43) |  |
| **Beck Depression Inventory** |  |  |  | 1.06E-15* |
| N-Miss | 1 | 0 | 1 |  |
| Mean (SD) | 4.3 (5.1) | 18.9 (9.7) | 11.5 (10.6) |  |
| Median (range) | 2.5 (0 - 21) | 18 (0 - 46) | 9 (0 - 46) |  |

Five years after enrollment, FM patients were contacted by telephone and asked about their outcomes using the Patient Global Impression of Change (PGIC) scale. Follow-up data were successfully obtained from 82% of the patients. Based on PGIC responses, participants were classified as Improvers or Persisters. The transcripomic profiles obtained at baseline were then retrospectively analyzed according to this 5-year outcome classification (see Materials and Methods for details).

### NEUTROPHIL PROFILING UNVEILS INFLAMMATORY LANDSCAPE IN FM PATIENTS

A transcriptome-wide comparison was first conducted between neutrophils collected from FM patients and those from HC (Figure. 1B). This analysis identified 56 genes with significant differential expression at the FDR 5% level (Table S1). Among those, 53 genes were upregulated in FM patients while only 3 genes were upregulated in HC (Figures. 1C and 1D). Pathway analysis was performed to distinguish the biological relevance of differences in the gene expression. Unbiased analysis of all the genes differentially expressed using Fgsea showed upregulation of 72% of the pathways in FM patients and only 28% in HC (Figure. 1E) (Table S2). Further pathway analysis focusing on statistically significant differentially expressed genes using gProfiler revealed that in both methods, the “inflammatory response” and “response to lipopolysaccharide” pathways were significantly overrepresented in FM samples (Figure. 1F) (Table S3). These analyses support the notion of a pro-inflammatory state in FM, characterized by elevated inflammatory processes in circulating neutrophils.

### DECIPHERING INEFFECTIVE LPS RESPONSES IN PATIENTS WITH FM

Inflammatory response induction via LPS was used to objectively quantify the responsiveness of neutrophils in both FM patients and HC (Figure. 2A). The analysis of LPS-mediated neutrophil activation in HC and FM samples identified over 10,000 transcripts in each group with significantly changed expression (Figure. 2B, Tables S4 and S5). There was a robust difference, however, in the magnitude of the response to LPS, which was much greater in HC compared to FM (p-value and D value of p-values distribution difference in Kolmogorov-Smirnov (KS) test < 2.2e^-16^, D = 0.38, and p-value and D value of log2 fold changes distribution difference KS test < 3.3e^-4^, D = 0.02; Figure. 2C) despite the absence of a significant difference in expression of the LPS receptors, *TLR4* and *LY96,* in FM versus HC samples (Table S1). The disparities became further pronounced upon conducting pathway analyses, wherein both Fgsea and gProfiler methods revealed a considerably higher number of LPS-dependent differentially regulated pathways in HC compared to FM (Figure. 2B) (Tables S6 to S9). Similarly, a more thorough pathway analysis of the top 300 significantly differentially expressed genes showed that HC neutrophils displayed 292 significantly enriched pathways with “inflammatory response” and “response to lipopolysaccharide” emerging as the most significant (Figure. 2D). In contrast, FM neutrophils had only 23 significant pathways with predominant presence of metabolic processes within the top three pathways. The anticipated “response to lipopolysaccharide” pathway was absent among the enriched pathways in FM (Figure. 2E). These results suggested that although the neutrophils from both FM and HC respond to LPS stimulation, the LPS-driven responses in FM are blunted and dysregulated.

**Figure. 2.**
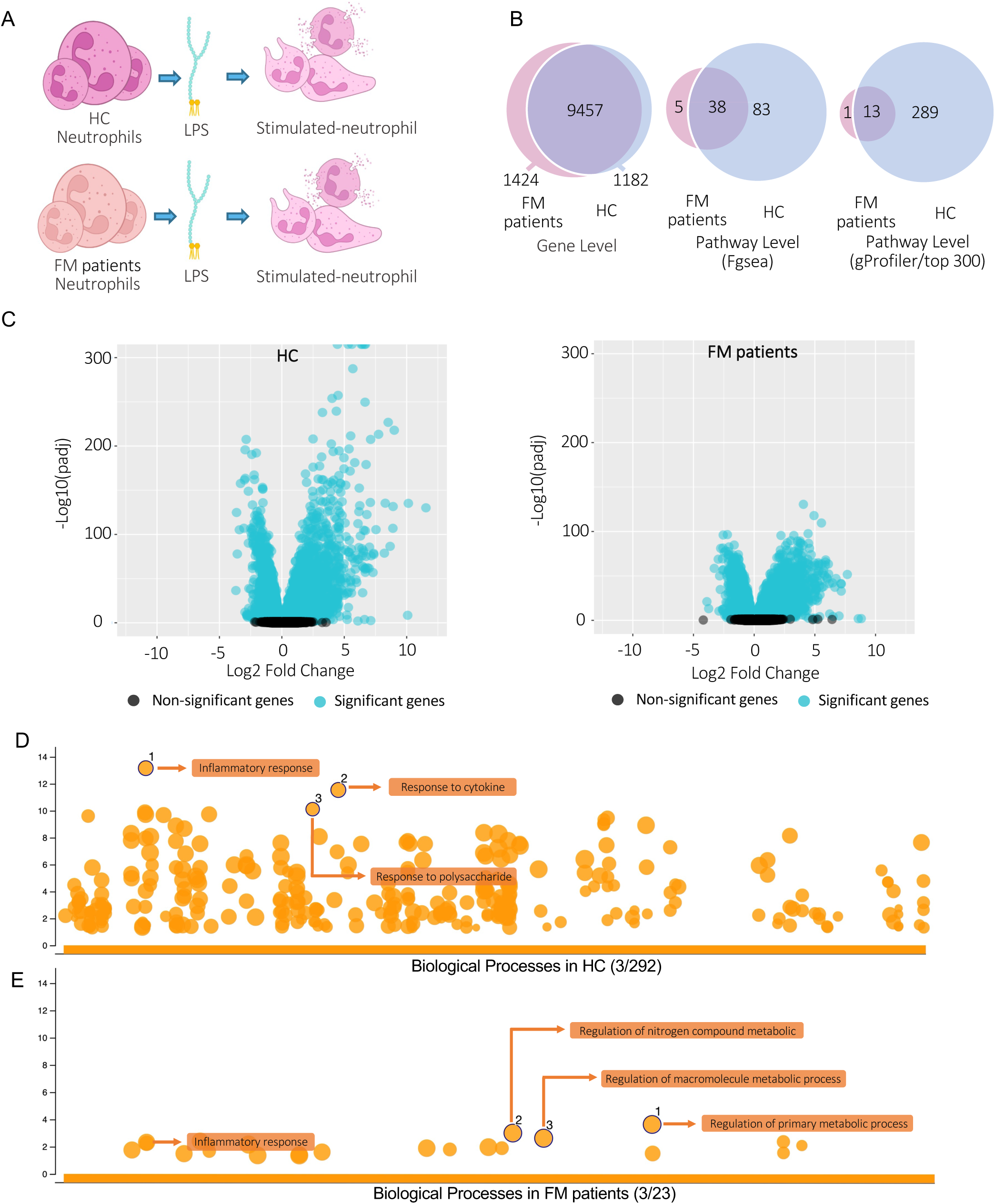
Neutrophils from FM patients show a blunted response to LPS stimulation. (A) Schematic representing the design of the experimental groups in the analysis. (B) Pie chart representing the counts of genes and pathways that are significantly differentially expressed after LPS stimulation. (C) Scatterplot representing the log2fold changes versus the statistical significance of the genes regulated by LPS stimulation. Black symbols represent genes regulated with an adjusted p-value of higher than 0.05, and blue symbols lower than p-value=0.05. (D) Bubble chart representing gProfiler pathway analysis results performed on the top 300 genes significantly regulated by LPS in HC. The x-axis represents the biological processes from GO. The y-axis shows the adjusted enrichment p-values in negative log10 scale. (E) Bubble chart representing gProfiler pathway analysis performed on the top 300 genes significantly regulated by LPS in FM patients. The x-axis represents the biological processes from GO. The y-axis shows the adjusted enrichment p-values in negative log10 scale.

### INTERACTION TERM ANALYSIS FURTHER ELUCIDATES DISTINCT DYNAMICS OF RESPONSIVENESS

Interaction term analysis was performed to identify genes with the largest difference in the trajectory of LPS stimulation of the neutrophils in FM patients compared to HC (Figure. 3A). Eighteen transcripts were identified as significantly differentially regulated within groups (Figure. 3B, Table S10). A consistent pattern was observed for 15 of these genes, wherein they exhibited upregulation in both FM patients and HC following stimulation with LPS (Figure. 3B). Notably, for all of these genes, despite the higher initial level of expression in FM patients compared to HC, they were incapable of reaching the maximal expression levels observed in HC following LPS stimulation. Importantly, the majority of these genes were well-known inflammatory markers, namely *TNF* and *IL6*, but also a few anti-inflammatory markers, namely *IL1RN* and *NFKBIA*. To better discern this expression pattern, the three key inflammatory markers are illustrated in Figure. 3C. These results further highlight that inflammatory-related genes are upregulated in non-stimulated neutrophils of FM patients but display a blunted response to LPS stimulation compared to HC. Of note, a few genes, including *DLGAP3* and *RYR3*, showed divergent response directions upon stimulation. We did not pursue these genes further, as our focus was on the more general pattern of regulation that was represented by inflammatory related genes.

**Figure. 3.**
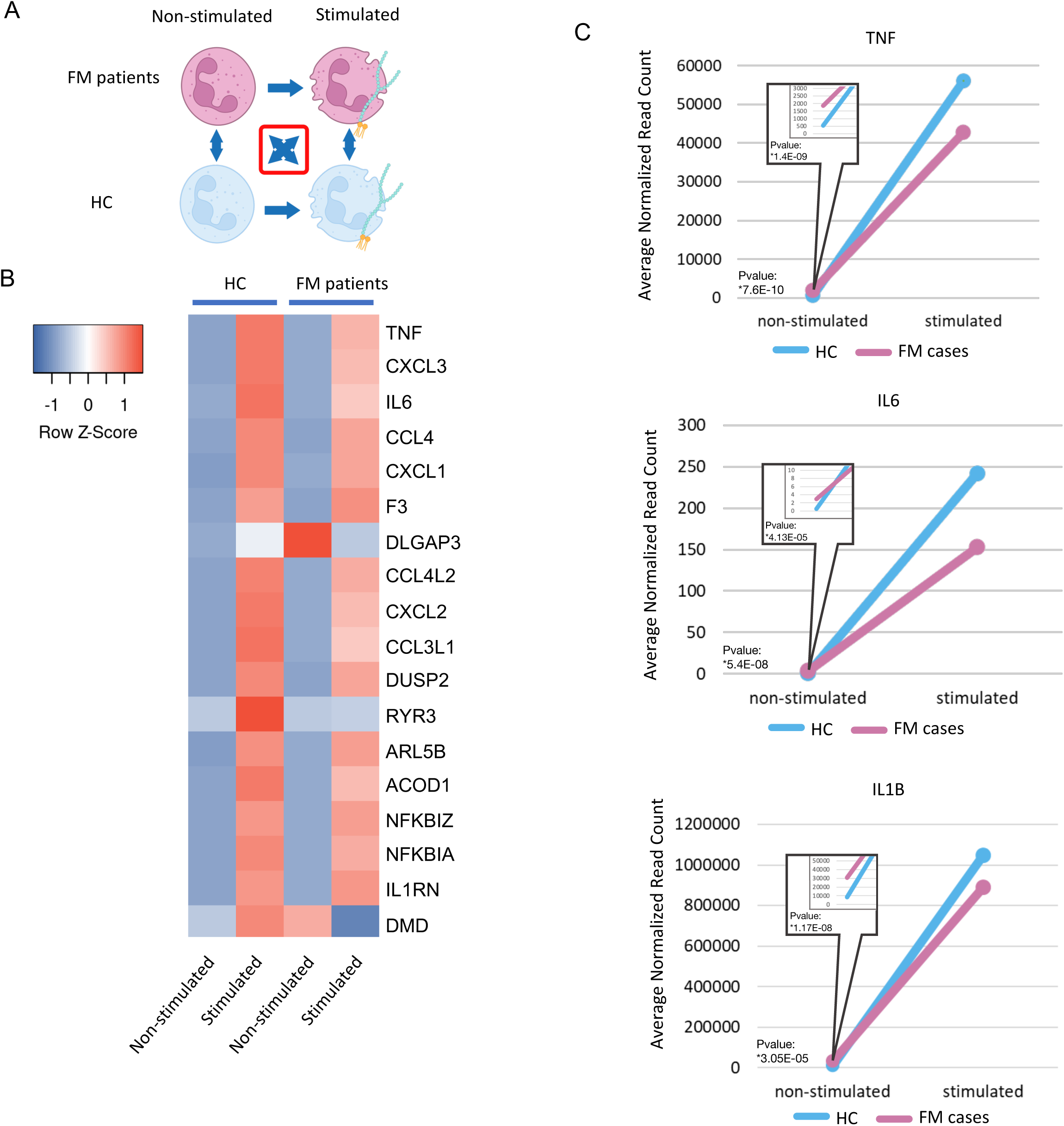
Interaction term analysis identifies genes responding differently to LPS stimulation in FM patients vs HC. (A) Schematic representing the design of the experimental groups for interaction term analysis. (B) Heatmap representing the genes that are significantly differentially expressed after LPS stimulation in FM patients versus HC considering pre- stimulation expression level. (C) Line graph representing gene expression patterns of three inflammatory markers in LPS-stimulated and non-stimulated conditions in FM patients and HC.

### LONG-TERM FM OUTCOME INTRODUCED TWO TRANSCRIPTIONALLY DIFFERENT SUBGROUPS OF PATIENTS

Using a patient-reported outcome of improvement obtained 5 years after enrollment, we divided FM patients into two subgroups, FM patients whose symptoms improved (Improvers, 41%, n=21) and FM patients whose symptoms did not improve (Persisters, 56%, n=30). We then studied the way neutrophils purified at enrollment responded to LPS and compared each subgroup to the HC group. We observed that the HC group has a much larger response to LPS than Improvers, and Improvers have a much larger response to LPS than Persisters (p-value and D value of p-values distribution difference in KS test < 2.2e-16, D = 0.28, and p-value and D value of log2 fold changes distribution difference in KS test < 2.2e-16, D = 0.11; Figure. 4A–C and Tables S11 and S12). Furthermore, a clearer dichotomy was identified between Improvers and Persisters when we compared the pre-stimulation state of each group with HC (Figure. 4D–F). While Persisters were the most different from HC (Figure. 4F), Improvers showed a substantial similarity to HC with no statistically different transcripts (Figure. 4E), suggesting that the difference in gene expression observed between FM patients and HC (Figure. 4D) is driven by Persisters.

**Figure. 4.**
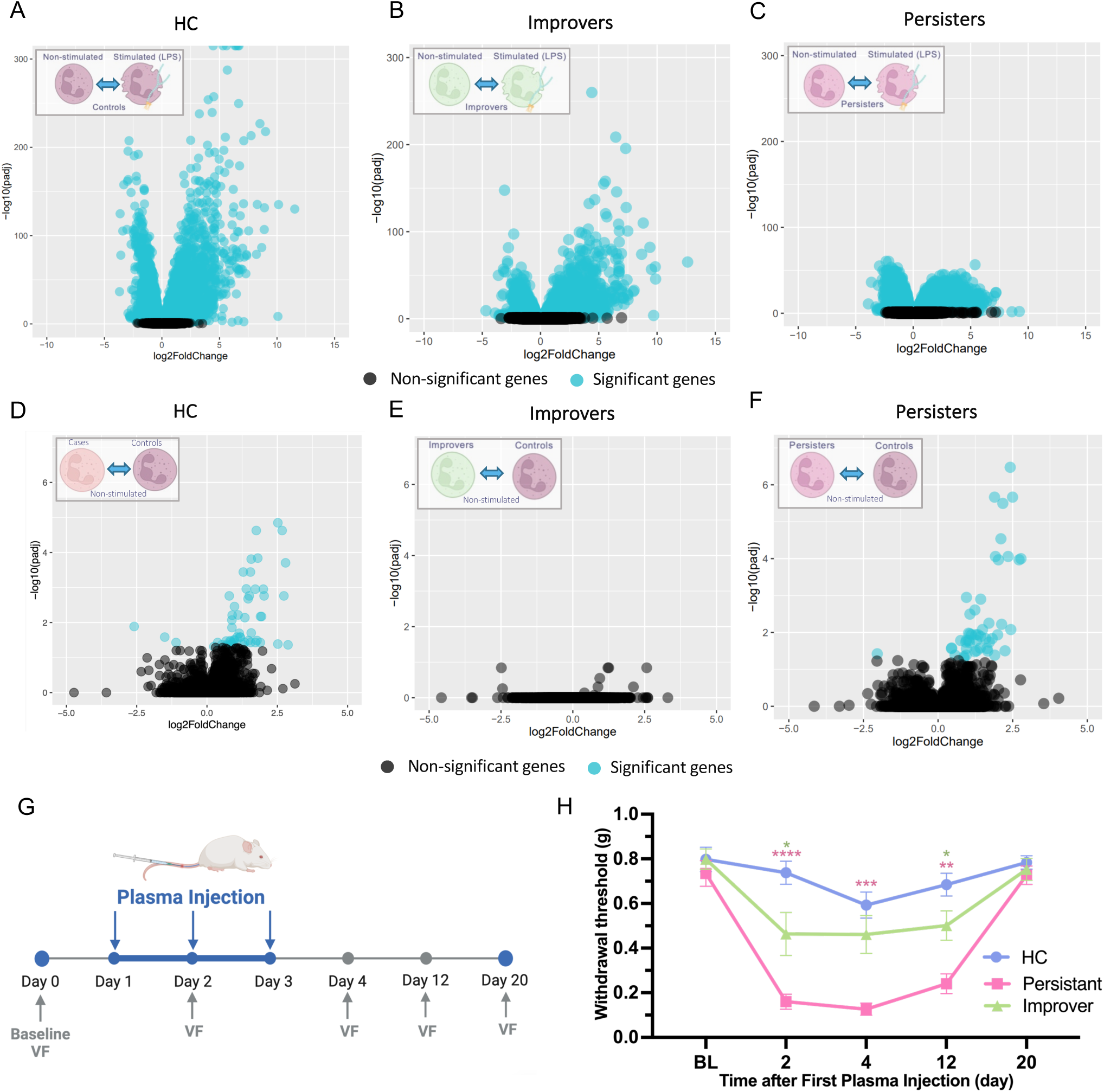
Neutrophils of FM Improvers and Persisters show a gradation in transcriptomics pattern. (A-C) Scatterplot representing the log2fold changes versus the statistical significance of the genes regulated by LPS stimulation in (A) HC, (B) FM Improvers, and (C) FM Persisters. (D- F) Scatterplot representing the log2fold changes versus the statistical significance of the genes differentially expressed in non-stimulated neutrophils when comparing HC and (D) FM patients, (E) Improvers, and (F) Persisters. Black symbols represent genes regulated with an adjusted p- value of higher than 0.05, and blue symbols lower than p-value=0.05. (G) Schematics representing the experimental design for FM plasma-dependent hypersensitivity in the mouse model. (H) Line graph representing mechanical paw-withdrawal thresholds before and after plasma injections from FM Persisters, FM Improvers, or HC. Symbols and bars represent mean ± SEM. *P < 0.05, **P < 0.01, and ***P< 0.001, compared to the corresponding HC (4 male and 4 female mice per group).

We then tested whether patient characteristics at enrollment were associated with long-term outcomes measured by PGIC. The analysis revealed a moderate negative correlation between PGIC and Fibromyalgia Survey Scores and PGIC (ρ = −0.44, p-value = 0.001), and Fibromyalgia Impact Questionnaire scores and (ρ = −0.44, p-value = 0.001). A weaker but significant negative correlation was also observed for the Symptom Checklist 90R (ρ = −0.32, p-value = 0.02). Additionally, lower BMI (ρ = −0.31, p-value = 0.03) and better sleep quality (PSQI: ρ = −0.30, p-value = 0.03) at baseline were associated with improved PGIC outcomes. No meaningful relationships were observed with age (ρ = 0.106, p-value = 0.46), sex (p-value = 0.81), pressure pain thresholds (ρ = 0.17, p-value = 0.25), perceived stress (ρ = 0.178, p-value = 0.21), anxiety (BAI: ρ = −0.233, p-value = 0.10), or depression (BDI: ρ = −0.265, p-value = 0.06) (Table 2).

**Table 2.** Correlation of clinical and demographic variables with Patient Global Impression of Change (PGIC) continuous scores. N represents the number of participants used in each correlation. P-values represent correlation test results (Spearman’s rank correlation results for continuous variables and point-biserial correlation for binary variables). A negative coefficient means a the lower the variable the higher the improvement. Asterisks (*) denote statistically significant results.

| Phenotype | Coefficient | # of subjects | p-value |
| --- | --- | --- | --- |
| Age | $\rho = 0.106$ | n = 51 | 0.46 |
| Sex (binary) | point-biserial = 0.24472 | n = 51 | 0.8077 |
| BMI | $\rho = -0.311$ | n = 51 | 0.0265* |
| Fibromyalgia Survey | $\rho = -0.44$ | n = 50 | 1.38E-03* |
| Beck Anxiety Inventory | $\rho = -0.233$ | n = 51 | 9.95E-02 |
| Fibromyalgia Impact Questionnaire | $\rho = -0.44$ | n = 50 | 1.40E-03* |
| Symptom Checklist 90 R | $\rho = -0.324$ | n = 51 | 2.04E-02* |
| Pressure Pain Threshold Score | $\rho = 0.17$ | n = 48 | 2.48E-01 |
| Pittsburgh Sleep Quality Index score | $\rho = -0.303$ | n = 50 | 3.25E-02* |
| Perceived Stress Scale | $\rho = 0.178$ | n = 51 | 0.213 |
| Beck Depression Inventory | $\rho = -0.265$ | n = 51 | 5.98E-02 |

Additionally, no identifiable significant associations were found between medication category or fibromyalgia duration with binary improvement status (Tables S13 and S14).

### PLASMA FROM FM PATIENTS CAUSED PROLONGED MECHANICAL HYPERSENSITIVITY IN MICE

To test if malfunctioning neutrophils from FM patients contribute to a pronociceptive environment, we injected pooled blood plasma collected at enrollment, from three randomly selected participants from each group, Improvers, Persisters, and HC, intravenously to naïve mice (Figure. 4G). Injection of FM patients’ plasma led to a significant decrease in mechanical paw-withdrawal thresholds at the 2-, 4- and 12-day time points in both groups (P<0.05). Notably, mice who received plasma from Persisters FM patients, had, however, a significantly greater decrease in mechanical paw withdrawal thresholds (P<0.001). All groups had recovered to baseline levels of mechanical threshold by day 20 post-transfer (Figure. 4H). These findings provide a new animal model of FM and substantiate the molecular distinctions among different FM patient subgroups.

### PLASMA FROM FM PATIENTS ACTIVATES SENSORY NEURONS

To test if plasma from FM patients could directly activate sensory neurons, we used an *in vitro* calcium imaging approach to measure evoked activity of cultured neurons in response to plasma derived from the same HC, Persister, and Improver patients. Representative images and calcium flux traces (Figure. 5A) demonstrate the capacity of human plasma to activate murine primary sensory neurons. Different patterns were chosen for each conditions but there was no pattern specific to the different plasma and all had neurons with the different patterns of response. Greater proportions of KCl-responding neurons responded to plasma from FM Improvers (35.8±1%) and Persisters (57.3±3%) compared to HC plasma (16.9±2%) (Figure. 5B; one-way ANOVA, P<0.001). Moreover, we observed a significant increase in the proportion of Na_v_1.8-negative cells responding to plasma from Improvers and Persisters compared to plasma from HC (Figure. 5C; one way ANOVA, P<0.05 for all comparisons except P>0.05 for Na_v_1.8+/ Nav1.8- Improvers vs Na_v_1.8+/ Na_v_1.8- Persisters proportions of responding cells). These data reflect the increased number of large-diameter neurons responding to FM Improvers or Persisters plasma (Figure. 5D). This phenotypic switch in response profiles of plasma-activated dorsal root ganglion (DRG) neurons suggests that plasma from FM Improvers and Persisters contains pro-nociceptive agents conferring the capacity to activate more heterogeneous populations of DRG neurons.

**Figure. 5.**
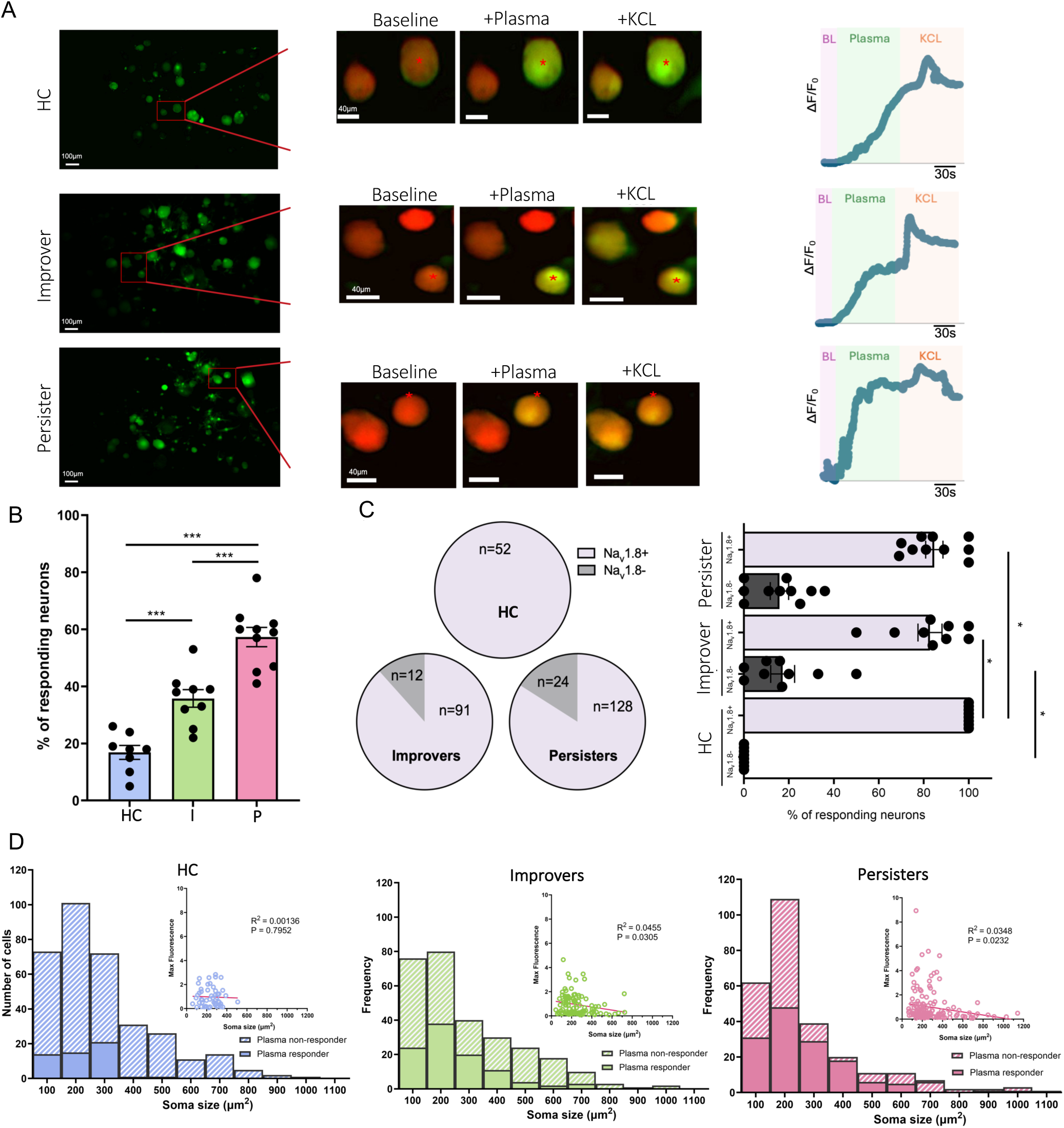
**Calcium imaging analysis of murine primary sensory neurons stimulated with human plasma from FM patients.** Plasma-evoked activity of primary sensory neurons in vitro. (A). DRG neurons in culture responding to plasma from HC (top panel), FM Improvers (middle panel), and FM Persisters (bottom panel). The magnified images with red stars over select cells indicate responding neurons with representative traces showing the change in calcium flux in the third column. (B) Bar chart representing the percent of neurons responding to plasma from HC, FM Improvers, and FM Persisters (***P<0.001 one way ANOVA). (C) Venn diagrams (left) and bar chart (right) representing the proportions of Nav1.8+ and Nav1.8- neurons responding to plasma from HC, FM Improvers, and FM Persisters (*P<0.05 one-way ANOVA). (D) Bar graphs representing the soma size distribution of responding and non-responding neurons and correlations of cell size with maximum fluorescence in cells responding to plasma from HC (blue), Improvers (green), and Persisters (pink).

### TRANSCRIPTION FACTOR ANALYSIS SUGGESTED NF-κB AS THE MULTI-LEVEL PRIMARY REGULATOR

To elucidate potential common regulatory factors among genes exhibiting differential expression between FM and HC at both non-stimulated and LPS-stimulated neutrophil states, TF analysis was conducted. The results of gProfiler based on TRANSFAC database indicated that genes differentially expressed in the non-stimulated state were orchestrated by the NF-κB complex (Figure. 6A) (Table S15). Interestingly, genes exhibiting differential responses to LPS and significant in the interactive model when comparing FM and HC were also predominantly regulated by different members of NF-κB transcription factor family (Figures. 6B and C and Table S16), which is only expected for the stimulated state as the LPS is a potent activator of NF-κB. TF analysis Enrichr of the same list of genes using TF protein–protein interactions (PPIs) database also identified NFKB1 as the only and most significant transcription factor both non-stimulated state (Adjusted p-value of 0.03) and the interaction term analysis genes (Adjusted p-value of 3.8e-5). To track the directionality of NF-κB activity at the pathway level, NF-κB-related pathways were extracted from the Fgsea results of differentially expressed pathways. We found that non-stimulated FM patients’ neutrophils expressed a higher level of “negative regulation of NF-κB transcription factor activity” pathway in comparison with HC. Within FM patients as well, this pathway showed higher levels in Persisters than in Improvers (Figure. 6D).

**Figure. 6.**
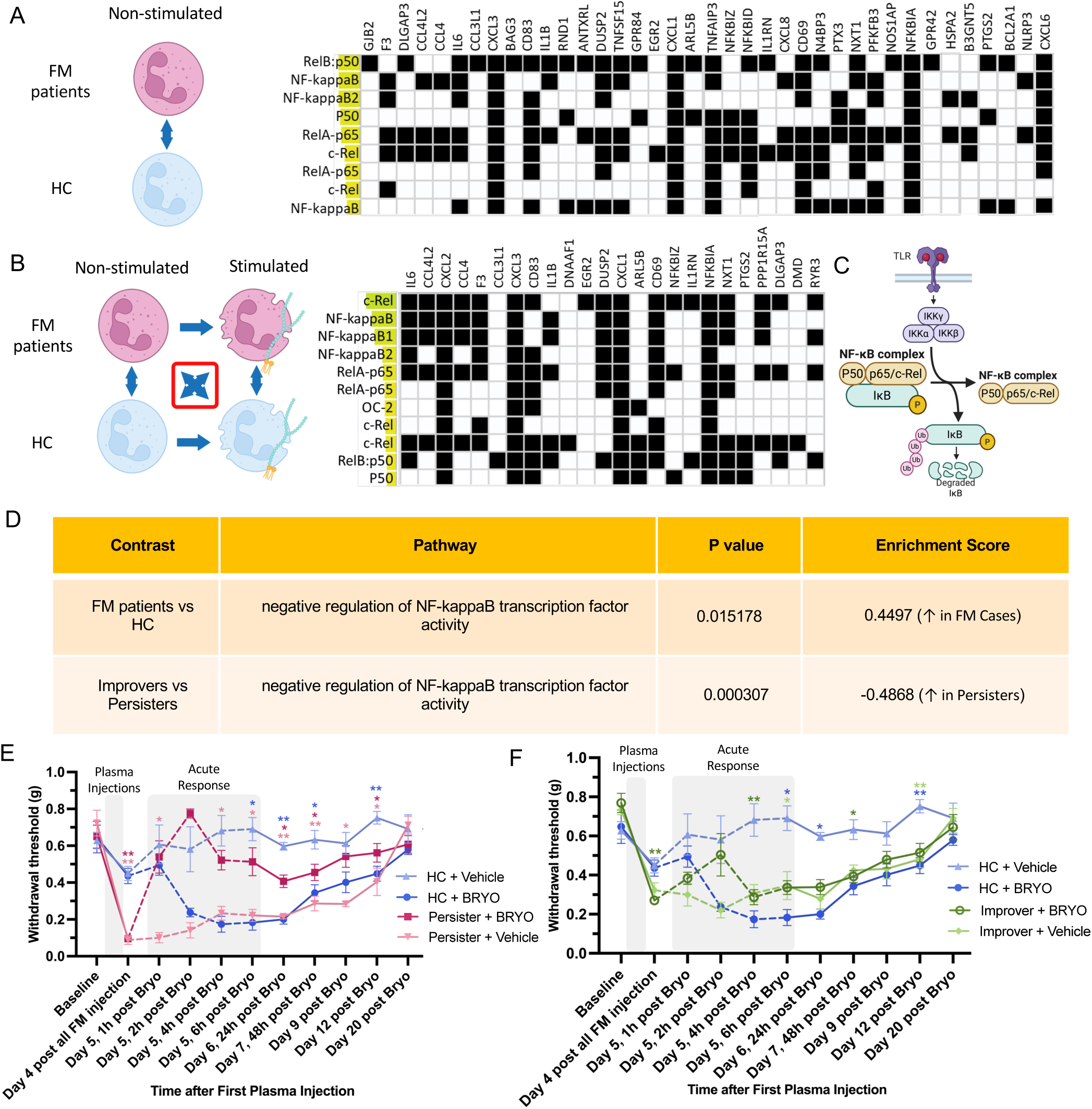
Significant NF-κB inhibition in FM patients points to a potential therapeutic target. (A-B) Schematic representing the design of the experimental groups and results for transcription factor analysis of differentially expressed genes at (A) comparison between non-stimulated neutrophils from FM patients and HC, and (B) interaction term analysis in LPS- stimulated and non-stimulated neutrophils from FM patients and HC. The X-axis represents the genes significantly contributing to the network of each TF; the Y-axis represents the names of the transcription factors. (C) Schematic representing the main components of the NF-κB complex. (D) NF-κB related pathways from pathway analysis performed for neutrophils from FM patients versus HC (first row), and FM Persisters versus Improvers (second row). (E-F) Line graphs representing Mechanical paw-withdrawal thresholds before and after injections of plasma from FM Persisters (E) or FM Improvers (F) versus HC, followed by injection of bryostatin or vehicle. Symbols represent mean ± SEM. *P < 0.05, **P < 0.01, and ***P < 0.001, compared to the corresponding HC + vehicle group (3 male and 3 female mice per group).

Because one of the main endogenous negative regulators of NF-κB, *NFKBIA* (NF-κB inhibitor alpha), was among the genes significantly regulated in our interaction term analysis (Figure. 3B), and its expression levels were positively associated with the expression of inflammatory genes before stimulation (Figure. S1), we conducted statistical tests to investigate whether *NFKBIA* is the upstream regulator of the expression levels of other differentially expressed genes. Instrumental variable (IV) regression analysis based on the pre-stimulated neutrophils indicated that changes in *NFKBIA* levels precede alterations in the expression levels of other genes, including *IL1B*, *CXCL3*, *CCL4*, *CXCL1*, *CCL4L2*, *ACOD1*, and *CXCL2*, mediated through IKKɑ (*CHUK*) (Figures. S2A–C). The Granger causality test that takes into account also LPS-stimulated outcomes further supported these findings, showing strong predictive evidence that *NFKBIA* regulates the expression of other genes (Figures. S2D and E). Notably, *NFKBIA* exhibited robust causal associations with all inflammatory genes, with only two non-inflammation-related genes failing to reach statistical significance. Together, our results suggest that the NF-κB transcription factor is the main source of impaired transcriptional regulation in the neutrophils of FM patients, such that NF-κB inhibition is associated with FM and disease persistence.

### NF-κB ACTIVATION REVERTS MECHANICAL HYPERSENSITIVITY INDUCED BY FM PLASMA INJECTION

Because our data suggested that impaired NF-κB function in neutrophils is associated with FM and its persistence, we hypothesized that reactivation of NF-κB may provide an avenue for alleviating FM-related pain. To test this hypothesis, we used a clinically available PKC-dependent activator of NF-κB, bryostatin, in our newly established animal model of FM plasma-transferred pain hypersensitivity. A single injection of bryostatin blocked mechanical hypersensitivity originating from FM Persisters plasma injection (p-value <0.05, compared to persistent + Veh) (Figure. 6E). However, in mice who received plasma from HC, as expected, bryostatin led to a significant decrease in mechanical thresholds, starting at 4 h and remaining for up to 96 h (Day 9) post injection (p-value <0.05, compared to HC + vehicle). In mice that received FM Improvers plasma, no significant difference was observed with the injection of bryostatin versus vehicle, although the same trend for improvement of hypersensitivity as in FM Persisters was observed (Figure. 6F).

### SINGLE-CELL PROFILING REVEALED INFLAMMATORY ACTIVATION ACROSS FM NEUTROPHIL SUBTYPES

To validate the presence of inflammatory processes in an independent FM cohort and to determine whether they are unique for neutrophils, we performed scRNA-seq of leukocytes of whole blood from three newly recruited FM patients and three HC, resulting in a total of 54,625 high-quality sequenced cells. Initial analysis comparing all blood cells between FM patients and HC revealed an elevated immune response and specifically an inflammatory response and response to cytokines in FM samples at both the gene and pathway levels, consistent with findings from bulk neutrophils’ RNA-seq (Figure. S3A–C, Tables S17 and S18). Focusing on neutrophils, we identified 25,743 high-quality neutrophil cells, which clustered into seven distinct groups (Figure. 7A). The distribution of neutrophils across clusters did not differ significantly between FM and HC (Figure. S3D), indicating the absence of evidence for an FM-specific neutrophil subtype. Differential gene expression analysis comparing neutrophils from FM and HC revealed a predominance of upregulated genes in FM samples (Figure. 7B, Table S19), which was not unique for neutrophils but was observed in the majority of blood cells (Figure. S4A–M). Enrichment analysis highlighted again an elevated innate immune, inflammatory response and response to cytokines in FM neutrophils, further supporting the dysregulated innate immune function observed in the bulk RNA-seq results (Figure. 7C, Table S20). Comparison of each of the cell subtypes in FM versus HC revealed a generally higher cellular activity in neutrophils of FM (Figure. S5A, Tables S21 and S22) with neutrophil clusters 7, 5, and 3 having the highest ratio of upregulated genes over all the significant regulated genes (Figure. S5B), although none of the neutrophil clusters were uniqly associated with FM. Consistent with our results in the discovery FM cohort, the *NFKBIA* gene was significantly upregulated in FM patients’ neutrophils in our scRNA-seq samples (Figures. 4B, 7D, and E, Tables S1, S10, S15 and S17). Notably, 4 out of 7 neutrophil clusters exhibited significant upregulation of *NFKBIA* in at least half of the cells within each cluster. Among them, Neutrophil Cluster 5 demonstrated the most pronounced upregulation, showing the highest statistical significance and the greatest log2 fold change (Figure. 7E). Additionally, TF analysis using the PPIs database replicated significant enrichment for *NFKB1* both at the level of whole blood (Adjusted p-value = 0.005) and the neutrophils (Adjusted p-value = 0.007).

**Figure. 7.**
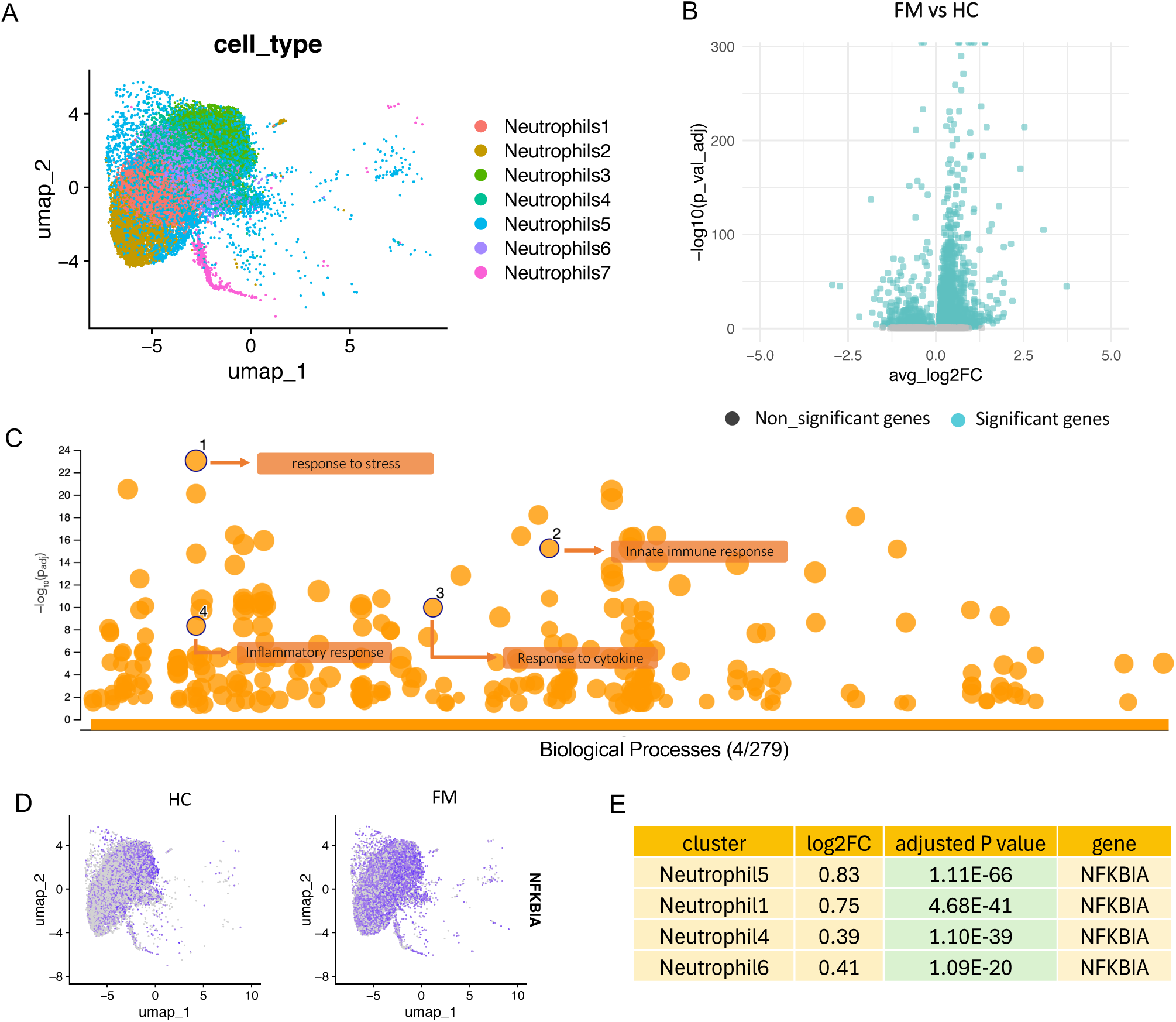
scRNA-seq shows an elevated innate immune response pattern in neutrophils of independent FM samples. (A) UMAP representing all identified neutrophil clusters. (B) Scatterplot representing the log2fold changes versus the statistical significance of the DEGs between FM and HC across all neutrophils’ clusters. Black symbols represent genes regulated with an adjusted p-value of higher than 0.05, and blue symbols represent p-value lower/equal to 0.05. (C) Bubble chart representing gProfiler pathway analysis results performed on the significant DEGs in neutrophils from FM patients and HC. The x-axis represents the biological processes from GO. The y-axis shows the adjusted enrichment p-values in negative log10 scale. (D) UMAP representing *NFKBIA* gene expression in FM and HC (adjusted p-value < 8.80E-302). (E) Differential gene expression results for the *NFKBIA* gene comparing FM versus HC in each neutrophil cluster separately. Positive log2 fold change indicates higher expression in FM patients.

## DISCUSSION

A large number of studies suggest the presence of low-grade inflammation in chronic pain patients, especially in FM patients (16, 20–22). This low-grade inflammation was also consisent with recent findings on DRG-invading neutrophils isolated from FM patients (*18*). Our initial analysis of neutrophils revealed a distinct inflammatory state in the neutrophils of FM patients compared to those of HC, in line with previous research (*23, 24*). However, recent evidence (*19*) also points to the unique role of a robust inflammatory response of short duration, driven by neutrophils, in promoting tissue healing (*25, 26*) and pain resolution and preventing the transition from acute to chronic pain in patients with acute low back pain and painful temporomandibular disorder. We thus hypothesized that neutrophils’ intrinsic potential to activate such inflammatory response might both prevent chronic pain development and resolve existing chronic pain. We sought to capture this potential for responsiveness in isolated neutrophils with an exogenous stimulus. We used LPS to evaluate the responsiveness of neutrophils which has served as a potent mediator of neutrophil activation in research for decades (27–29). Comparing FM patients and HC revealed a substantial difference in the magnitude of the LPS response between the clinical groups. Further investigations demonstrated that whereas the pre-stimulation expression of the majority of the significantly differentially expressed genes—including key inflammatory markers such as *CCL4*, *TNF*, and *IL6—*is higher in FM neutrophils, the magnitude of their increased expression after LPS stimulation is lower than that of HC. This occurs despite the absence of any differences in the expression of LPS receptors themselves (Table S1). Similarly, unlike healthy controls, where LPS stimulation prominently activates “inflammatory response” and “response to LPS” pathways, neutrophils from FM patients show minimal transcriptional changes following LPS exposure, with most enriched pathways limited to metabolic processes. These findings highlight a markedly inefficient immune activation in FM neutrophils.

We then discovered that the neutrophils’ response to an inflammatory stimulus along with clinical symptoms may serve as a reliable prognostic marker for FM persistence. Reanalysis of the data, incorporating long-term follow-up status, revealed two transcriptionally heterogeneous FM subgroups, defined in this study as Improvers and Persisters. Notably, these clinical sub-groups exhibited blunted levels of LPS response in comparison with HC, with a positive relationship between the strength of LPS response and a favourable long-term outcome. In contrast, only Persisters displayed a distinct inflammatory profile pre-stimulation by LPS compared to HC. This transcriptional heterogeneity was further validated *in vivo* and *in vitro* using blood plasma. Interestingly, in the animal assays, injection of plasma from FM patients induced robust mechanical hypersensitivity lasting for two weeks. This effect was of considerably greater magnitude in the Persisters group than in the Improvers group and completely absent in HC. We suggest that this assay should be more widely adopted as an animal model of FM pain, helping to create a painful immune state in animals with obvious translational relevance. Similarly, FM plasma, particularly from Persisters, activated a much higher number and a wider range of murine cultured sensory neurons, including Na_v_1.8-negative neurons, suggesting a phenotypic switch in sensory neurons activity that may involve increased expression of pro-nociceptive ion channels and altered neurotransmitter release, contributing to heightened neuronal excitability and sensitivity (*30, 31*). Together, these results suggest that plasma from FM patients contains pro-nociceptive agents possibly resulting, at least partially, from the malfunctioning neutrophils (*32*). Our investigation into the upstream regulator of the impaired cellular response in FM patients pointed to NF-κB, a major inflammatory activator, as the preeminent and most significant transcription factor. Unexpectedly, pathway analysis revealed that FM patients have a significant *negative* regulation of NF-κB transcription factor activity. Similarly, this inhibitory effect was more pronounced in the Persisters subgroup compared to the Improvers. Although inflammation, particularly that involving NF-κB activation, has been traditionally viewed as hyperalgesic (33–36), with many studies exploring NF-κB inhibition as a pain therapy, our results suggest a more nuanced role. Additionally, causal modelling using IV regression and Granger analysis identified NFKBIA, the major NF-κB inhibitor, as a central upstream and causal regulator of inflammatory gene expression in FM patients, including IL1B, CXCL1, and CCL4, and not just an outcome of more inflammation in FM patients. These findings provide strong support for NF-κB inhibition as a driver of the dysfunctional neutrophil response observed in FM.

We previously proposed that controlled reactivation of inflammation may be necessary for resolution and long-term pain relief (*19*), challenging the idea that pain can be effectively managed solely by suppressing inflammation. Many studies supporting NF-κB inhibition focus on short-term effects and overlook potential long-term consequences (37–39). Recent evidence suggests that activating, rather than suppressing, immune responses may promote sustained analgesia by triggering negative feedback loops that reduce inflammatory cytokine production (*40*). Excessive NF-κB inhibition likely impairs neutrophil response to inflammatory stimuli like LPS, leading to an incomplete inflammatory reaction and failure to initiate homeostatic feedback essential for pain resolution. In FM, low-grade inflammation may drive this compensatory NF-κB suppression, hindering recovery. To test this, we used the PKC-dependent NF-κB activator bryostatin in animal models (*33, 41*), in contrast to traditional inhibitory approaches, as a strategy to stimulate the inflammatory response. Bryostatin produced immediate antinociception and prevented FM plasma-induced hypersensitivity, with stronger effects in mice treated with plasma from Persisters than Improvers. Consistent with prior work, bryostatin induced mechanical hypersensitivity in mice receiving plasma from HC, reflecting expected NF-κB activation pro-nociceptive effects (*42*). These findings suggest that NF-κB activation via agents like bryostatin, a drug used for other indications (*43, 43, 44, 44*), may offer a novel therapeutic strategy for FM and potentially other chronic pain disorders.

Next, in order to replicate our results and investigate immune cell profiles in FM patients with high resolution, we performed, scRNA-seq on whole blood samples from FM patients and HC. Our results validated our findings from bulk RNA-seq related to the inflammatory phenotype of neutrophils from FM patients, and also revealed that while cellular inflammatory phenotype was not confined solely to neutrophils, it was most pronounced in neutrophils and monocytes (Table S22). Importantly, within the neutrophil population, although all subclusters exhibited a degree of elevated immune response (Tables S21 and S22), only four subclusters demonstrated high expression of *NFKBIA*, the major inhibitor of NF-κB. Among all the subclusters, *NFKBIA* had the highest expression in the Neutrophil5 cluster. This cluster also showed exclusive predominant expression of Fcγ receptors, *FCGR3B* and *FCGR3A*, which are typically associated with antibody-dependent cellular responses (*45, 46*), and their overexpression aligns with FM features like autoantibody prevalence and glial cell activation, potentially contributing to neuroimmune cross- talk and centralized pain mechanisms (18, 47–49)(Tables S23 and S24). Notably, these findings suggest that even though none of the neutrophil subclusters were uniqly associated with FM, these NFKBIA-high neutrophil subclusters, and especially Neutrophil5, may potentially contribute to the impairment of pain resolution in chronic pain patients through a diminished capacity to mount an adequate inflammatory response upon activation.

Previous studies have characterized functionally distinct neutrophil phenotypes, notably the pro-inflammatory N1 and the pro-tumoral, immunosuppressive N2 subsets, particularly within the context of cancer (50–52). However, in our analysis, we did not identify a discrete subpopulation of neutrophils that clearly aligned with the classical N1 or N2 phenotypic signatures. This indicates that neutrophil functional polarization in our dataset may not follow the traditional N1/N2 paradigm, but instead reflects a previously suggested transitional spectrum of states shaped by chronic pain (*53, 54*).

In conclusion, our investigation of peripheral blood neutrophils has uncovered a distinct inflammatory state in FM patients, but also an inefficient response to LPS, implicating a potential cell signalling defect leading to chronic, low-grade inflammation. NF-κB transcription factor inhibition emerged as a crucial regulator in this context. Moreover, our animal experiments supported the relevance of these findings, emphasizing the pivotal role of NF-κB activation in alleviating FM plasma-driven hypersensitivity. The identification of transcriptionally dependent gradation of FM sub-groups further contributes to our understanding of FM, lays the foundation for pathophysiologically based biomarkers of heterogeneous FM patients, and provides valuable insights for the development of new effective treatment strategies.

## MATERIALS AND METHODS

### SEX AS A BIOLOGICAL VARIABLE

Both male and female patients and controls were enrolled in this study. However, because fibromyalgia predominantly affects women, the number of male participants was small and unbalanced relative to female participants. As a result, direct sex-based comparisons were not feasible. Nevertheless, sex was included as a covariate in all analyses. In the animal experiments, both male and female mice were used, and no sex-related differences were observed.

### STUDY DESIGN AND SPECIMEN COLLECTION

To characterize the transcriptional profile of FM immune cells, a cohort of FM patients and HC were analyzed. The cohort was collected between August 5, 2016, and September 7, 2017 as described in detail in Verma et al. (*55*) In brief, fibromyalgia was diagnosed by the study physician in accordance with the 2010 diagnostic criteria delineated by the American College of Rheumatology (*56*). During the visit, blood was collected and neutrophils were purified from 62 FM patients and 64 matched HC. Fibromyalgia Survey, Fibromyalgia Impact Questionnaire (FIQ), Symptom Checklist 90 R (SCL-90-R), Pittsburgh Sleep Quality Index score (PSQI), Perceived Stress Scale (PSS), Beck Anxiety Inventory (BAI), and Beck Depression Inventory (BDI) questionnaires (57–61), among others, and testing of pressure pain thresholds (PPT) (*62*) were administrated to gather comprehensive clinical and psychological information. Medication data was also collected at the time of recruitment including both medications prescribed specifically for FM and those for other conditions.

To assess the duration of FM, an email survey was sent in December 2020 to all FM patients who had previously consented to be recontacted. Participants were asked about the year their pain began and the year they got the diagnosis. Additionally, to study long-term outcomes and their underlying mechanisms in FM patients, 45 participants who had provided recontact consent were questioned by phone interview between July and August of 2022. The Patient Global Impression of Change questionnaire (PGIC) (*63*) was employed to assess overall clinical condition change since initial study participation. During this process patients were asked, "Since your participation in our study how would you describe the change (if any) in ACTIVITY LIMITATIONS, SYMPTOMS, EMOTIONS, and OVERALL QUALITY OF LIFE, related to your painful condition?”. Based on the patients’ responses, FM patients were grouped as “Improver” or “Persister”. Participants who reported “A great deal better, and a considerable improvement that has made all the difference”,

“Better, and a definite improvement that has made a real and worthwhile difference”, or “Moderately better, and a slight but noticeable change” were defined as Improvers and those who reported “No change (or condition has got worse”, “Almost the same, hardly any change at all”, and “A little better, but no noticeable change” were defined as Persisters. Participants recruited for scRNA-seq were selected using the same criteria as the original cohort. All participants reported receiving a fibromyalgia diagnosis from a physician and were members of the Association de Fibromyalgie du Quebec. Our study examined male and female animals, and similar findings are reported for both sexes.

### PROCESSING OF BLOOD SAMPLES

Neutrophils were isolated from whole blood using magnetic labelling and separation technology as per manufacturer instructions (MACSxpress Whole Blood Neutrophil Isolation Kit, Miltenyi Biotec). Isolated neutrophils were counted (Cellometer Auto T4, Nexcelom) and resuspended at 1 million cells/mL in RPMI 1640 complete medium (Lonza) supplemented with 2 mM L-Glutamine, 10% fetal bovine serum, and 1% penicillin/streptomycin (neutrophil medium). Cells were then plated in 24-well plates (1 mL/well). One hundred μL of lipopolysaccharide (LPS) diluted in neutrophil medium (110 ng/mL) was added to half of the plated wells (final LPS concentration: 10 ng/mL) to activate neutrophils. The other wells received 100 μL of neutrophil medium (non-activated neutrophils). Following 1 h of incubation (37 °C, 5% CO_2_), cells were gently pipetted into a 15-mL conical tube and spun down at 300 *g* for 5 min at room temperature. Pelleted cells were resuspended in 650 μL of RLT buffer (QIAGEN) supplemented with β-mercaptoethanol (10 μL of 14.5 M β-ME/ 1 mL of RLT buffer) to support the inactivation of RNases and stored at -80 °C until RNA extraction. RNA was extracted using the AllPrep DNA/RNA Mini Kit from Qiagen.

The samples were sequenced at Genome Quebec Center in Montreal using Illumina NovaSeq 6000 with 30M depth. Plasma samples were obtained from 2 mL of whole blood, by centrifugation at 1200 *g* for 10 min and kept in -80 °C for later use. For scRNA-seq of whole blood, fresh samples were processed immediately. Red blood cell lysis was performed using a 1X ammonium chloride-based lysis buffer (10X stock: 8.02 g NH₄Cl, 0.84 g NaHCO₃, 0.37 g EDTA, QS to 100 mL with Millipore water). Whole blood was incubated with 10 mL of 1X RBC lysis buffer per 1 mL of blood at room temperature for 10–15 minutes until the sample became clear, indicating complete lysis. The lysed samples were centrifuged at 500 × g for 5 minutes at room temperature, and the supernatant was removed. The resulting cell pellet was used for cell capture and library preparation using HIVE CLX kits by McGill Genome Innovating Center, following the manufacturer’s instructions.

### ANIMAL EXPERIMENTS

CD-1 male and female mice aged 6–10 weeks (ICR: Crl, Charles River, St. Constant, QC) were used in these experiments. All mice were housed in standard shoebox cages with 2–4 (same-sex) per cage in a light (14:10 h, lights on at 07:00 h) and temperature-controlled (20±1 °C) environment with *ad libitum* access to food (Harlan Teklad 8604) and tap water. Mice were acclimated to the vivarium for 7 days post-arrival and before testing. Each mouse was used in a single experiment.

### PAIN BEHAVIOR ASSAYS

To analyze mouse pain behaviour, the mechanical paw-withdrawal threshold was measured with von Frey filaments using the up-down staircase method of Dixon (*64*). In every experiment, a baseline threshold determination was made, followed by reassessments post-plasma transfer and/or bryostatin injection at regular intervals (hours and/or days). All testing was performed during the light phase at approximately the same time each day. Prior to testing, mice were habituated in cubicles for about 2 h.

### EXPERIMENTAL ASSAYS

To assess the effect of plasma derived from two sub-groups of FM patients on mouse pain behavior, mice received three intravenous injections (tail vein, 0.2 ml)) of human plasma pooled from from 5 FM patients who improved their pain symptoms (Improvers) over the 5–6 years of follow up, 5 FM patients who had their pain symptoms persist (Persisters), or 5 HC. Mechanical paw-withdrawal threshold was assessed at baseline and days 2, 4, 12 and 20 post-plasma injection. To assess the role of NF-κB on the mechanical hypersensitivity induced by FM plasma injections, a second experiment was performed in which four days after plasma injections (of Improvers, Persisters, or HC), mice received an intraperitoneal injection of the NF-κB activator, bryostatin-1 (20 µl/ml, 0.2 ml in 1% DMSO, Tocris Bioscience, Bristol, UK) or vehicle (1% DMSO). Mechanical paw-withdrawal threshold was assessed at baseline, day 4 post-plasma injection (prior to bryostatin-1) and 1, 2, 4, 24, 48 h and 9, 12, and 20 days post-injection of bryostatin (on day 5).

### PRIMARY CELL CULTURE

Primary DRG neurons were isolated from male mice generated by crossing Na_v_1.8-Cre mice with TdTomato stop-floxed mice on a C57BL/6J background. Briefly, all ganglia were dissected from the entire length of the spinal column and transferred to a preequilibrated solution of Hanks’ balanced salt solution (HBSS) containing collagenase (type XI; 5 mg/ml), dispase (10 mg/ml), Hepes (5 mM), and glucose (10 mM) for 30 min (37 °C, 5% CO_2_). Ganglia were then gently centrifuged at 300 rpm for 5 min, and the supernatant was discarded and replaced with warmed Dulbecco’s modified Eagle’s medium (DMEM) GlutaMAX supplemented with 10% fetal bovine serum (FBS). The DRGs were mechanically triturated using a 1-ml pipette tip. The dissociated cells were then centrifuged at 300 rpm for 8 min, and the supernatant was removed and replaced with an appropriate volume of DMEM GlutaMAX supplemented with 10% FBS and nerve growth factors (50 ng/ml). Cells were plated onto sterilized glass coverslips (13 mm, size 0), precoated with poly-l-lysine (1 mg/ml) and laminin (1 mg/ml) for 0.5–2 h and then topped up with 2 ml DMEM GlutaMAX supplemented with NGF. Cells were maintained at 37 °C (5% CO_2_) for 16– 24 h before imaging.

### CALCIUM IMAGING

DRG cells were incubated in well plates with Fluo-4 (10 uM) diluted in extracellular solution (140 mM NaCl, 4 mM KCl, 1 mM MgCl2, 1.8 mM CaCl_2_, 10 mM Hepes, 5 mM glucose (pH 7.4 with NaOH)) for 30 min. Coverslipped DRG well plates were then placed into a perfusion chamber attached to an inverted microscope (Leica fluorescent DM IL LED Fluo microscope) and imaged using a 20X objective. During each imaging experiment, the neurons were continuously perfused using a gravity-fed perfusion system and images (488 filter) were recorded every 0.35 sec for a total of 4.5 min in response to the following stimulations: 30 sec standard solution, 2 min HC/Improvers/Persisters plasma diluted in standard solution (100 µl of plasma diluted in 10 ml standard solution with 33 µl per coverslip), 2 min 70 mM KCl solution (74 mM NaCl, 70 mM KCl, 1 mM MgCl2, 1.8 mM CaCl_2_, 10 mM Hepes, 5 mM glucose (pH 7.4 with NaOH)). Recorded TIFF videos were first stabilized using the MOCO ImageJ plugin, and ROIs were manually drawn within the cytoplasm of identified neurons. Raw data was extracted in the form of averaged pixels per ROI per frame and change in fluorescence over time (ΔF/Δt) calculated per frame. A neuron was classified as a responder to plasma if the following condition was met: the rate of change of stimulus-induced force (ΔFstim/Δt) is greater than the sum of the rate of change of basal force (ΔFbasal/Δt) and four times the basal force (4sbasal), where Fstim represents the most significant value of the derivative during a specific period of stimulus application. Fbasal represents the mean derivative baseline value, calculated by averaging the derivative values of the four frames preceding the stimulus. Sbasal, on the other hand, refers to the standard deviation of the baseline derivative values. The peak fluorescence of a specific neuron was found by calculating the change in fluorescence over the baseline, represented as ΔF/F0 in smoothed traces. ΔF/F0 is the difference between the fluorescence at a given time (Ft) and the minimum fluorescence observed during the baseline and stimulation period (F0), divided by F0.

### STATISTICS

The transcriptomics data were aligned to the human genome GRCh37/hg19 using the STAR aligner (*65*). Differential gene expression and interaction term analyses were performed using moderated statistical tests implemented in the R statistical package DESeq2 (*66*). Age, sex, and BMI were used as co-variates throughout the analysis. Pathway analyses were performed using “Fgsea” with statistical significance assessed based on an adaptive multi-level split Monte-Carlo scheme (*67*) as well as the gProfiler R package (*68*). Transcription factor analysis was performed using the gProfiler R package which takes its putative transcription factor binding sites from the TRANSFAC database (*68, 69*), as well as Enrichr using the PPIs database (*70, 71*). Transcription factor analysis was limited to the top-ranking genes differentially expressed between FM patients and HC at 5% FDR (56 genes) and in the interaction term results at 10% FDR to get a fair number of genes (27 genes). Adjustment for multiple comparisons was done using the False Discovery Rate (FDR) method. A significance threshold of 5% FDR was used. The causal relationship between *NFKBIA* and various target genes significant in the interaction term analysis was assessed in the pre-stimulation gene expressions derived from isolated neutrophils’ bulk RNA-seq data using instrumental variable (IV) regression analysis with the *CHUK* gene as an instrument. The analysis was performed in R using the ivreg function from the AER package. Fifteen genes out of 18 had calculated TPM and were available for this analysis. The statistical significance of each model was assessed using standard regression diagnostics, including tests for weak instruments. Granger causality tests were done using the grangertest function from the lmtest R package (*72*) to assess whether changes in *NFKBIA* expression predict alterations in inflammatory gene expression. The time-series data were constructed using a combined dataset that included both stimulated and non-stimulated conditions. scRNA-seq reads were mapped using BeeNet v2.X.X from Honeycomb Technologies (*73*). Data processing and analysis were conducted in R, utilizing the Seurat R package. Cell types were annotated using ScType (*74*). In order to ensure that only robust and biologically relevant genes were considered, we applied a minimum percentage threshold (min.pct = 0.5), requiring genes to be expressed in at least 50% of cells in either group for all the differential gene expression analyses.

Group comparisons of demographic and clinical variables between FM and HC were performed using Mann-Whitney U tests for continuous variables and Fisher’s exact test for categorical variables. Associations between baseline clinical characteristics and long-term outcomes measured by PGIC were assessed. The PGIC consisted of seven response options, coded numerically from 1 (“no change or condition has got worse”) to 7 (“a great deal better and a considerable improvement that made all the difference”). Continuous baseline measures were correlated with PGIC scores using Spearman’s rank correlation, chosen to avoid assuming equal spacing between response categories. Categorical baseline variables were correlated with PGIC score using point-biserial correlation. To evaluate the association between medication use and diagnostic group, Fisher’s Exact Test was used. The relationship between FM duration and self-reported improvement was assessed using Welch’s t-test and the Wilcoxon rank-sum test. All statistical tests were two-tailed, and significance was defined as p-value < 0.05.

### STUDY APPROVAL

The human cohort was collected under a research project approved by the Institutional Review Board (IRB) of the Faculty of Medicine, McGill University (Study Number A05-M50-14B). All animal procedures were approved by the Downtown Animal Care Committee at McGill University (Protocol Number MCGL-10077).

## DATA AVAILABILITY

All data associated with this study are present in the paper or the Supplementary Materials. The sequencing data is available upon request.

## AUTHOR CONTRIBUTIONS

Conceptualization: LD, SJE, CBM

Methodology: LD, SJE, MP, AO, LVL, SS, CBM, FM

Cohort collection: CBM, FM, JV Visualization: SJE, AO, SC Funding acquisition: LD, JSM, SS

Project administration: LD, SJE, FM Supervision: LD, JSM, SS

Analysis: SJE, MP, AO, LVL, JSM, SC, SS

Experimental Execution: AO, LVL, SC Writing – original draft: SJE, LD

Writing – review & editing: all co-authors

## FUNDING SUPPORT

Canadian Excellence Research Chairs funds (LD, CERC09) Canadian Institutes for Health Research Foundation Grant (JSM) Pfizer Canada Professorship in Pain Research (LD)

Canadian Institutes for Health Research Strategy for Patient-Oriented Research (SPOR) Orthopedic Research UK (558)

The Royal Society (RGS\R2\222387)

Edwards PhD. Studentships in Pain Research (SJE)

## ACKNOWLEDGMENTS

We would like to thank all the participants enrolled in this study, as without their contribution, this study would not have been possible. The schematic diagrams were created using BioRender.com (*BioRender.com/i91d301*). We thank McGill Genome Centre for sample processing and sequencing. We also acknowledge the support of our funders.

---

**Figure. S1.**
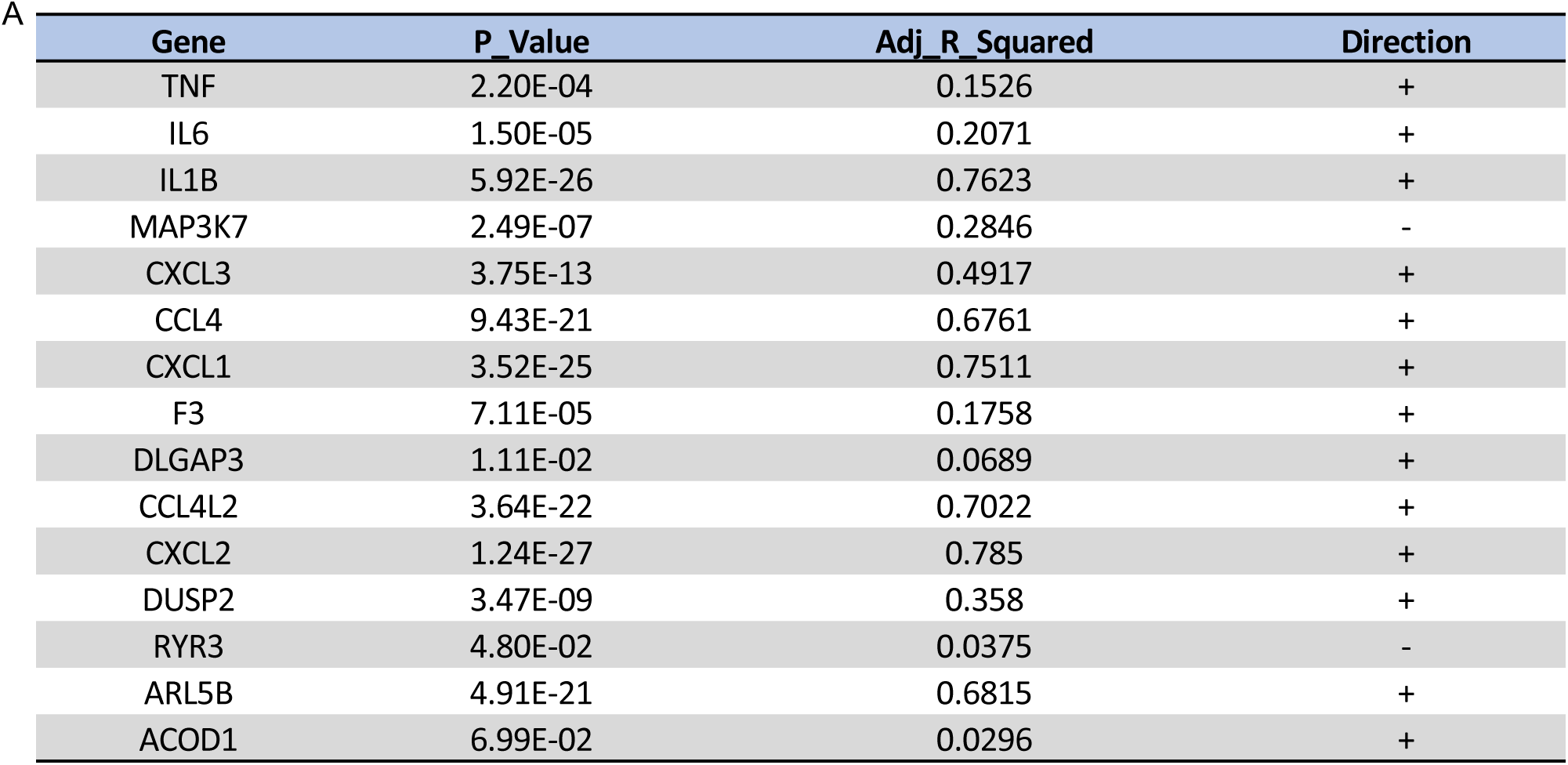
*NFKBIA* expression is positively associated with the expression of inflammatory genes in non-stimulated neutrophils in FM patients. Linear regression analysis of pre- stimulation gene expression with NFKBIA. The table shows p-values, adjusted R², and direction of association for each gene.

**Figure. S2.**
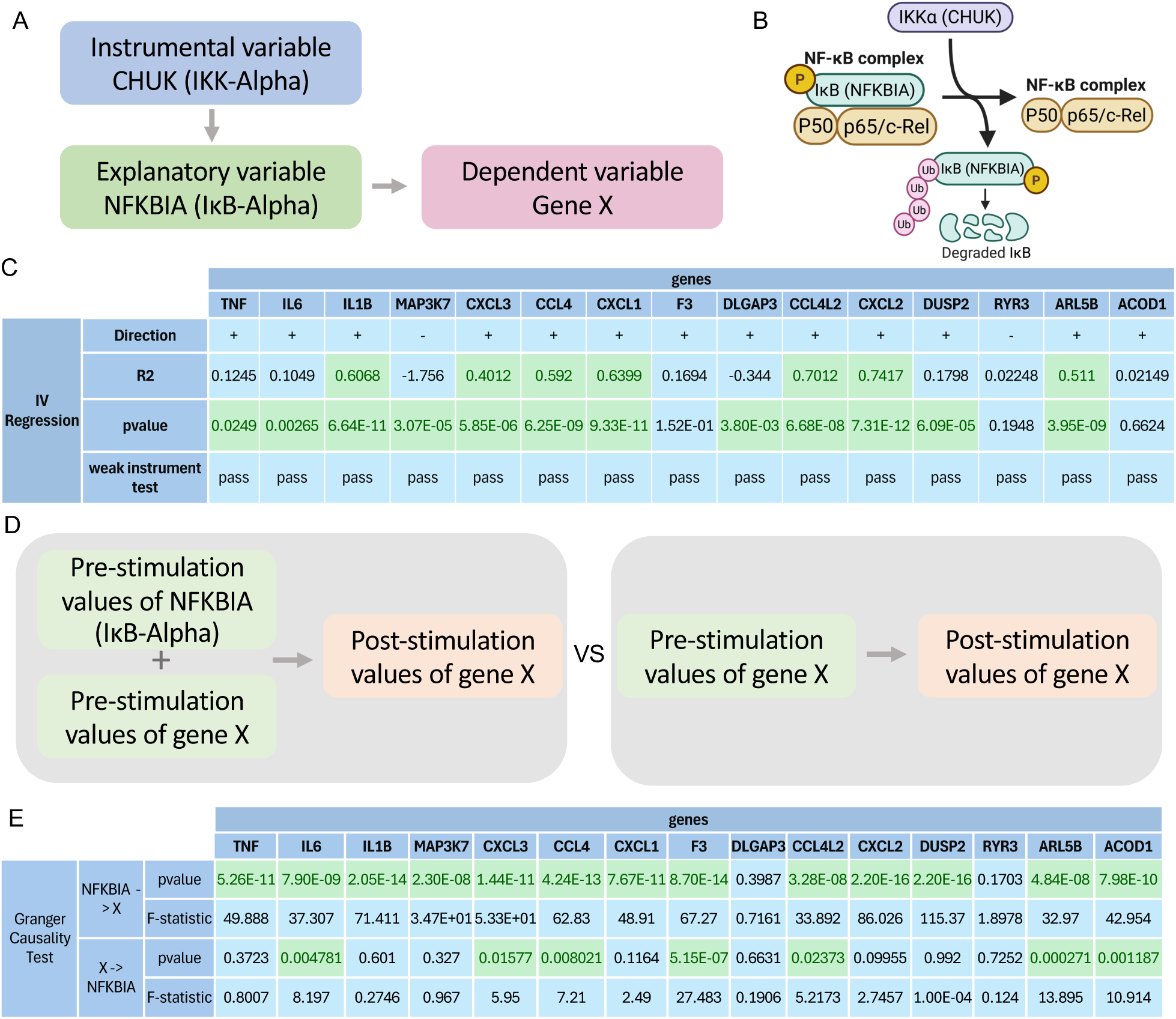
The Level of *NFKBIA* expression is causally related to the expression of inflammatory genes in FM. The causality was tested between expression levels of *NFKBIA* and the 15 genes significant in interaction term analysis (Figure.3B). (A) The diagram representing Instrumental Variable IV regression test illustrates the hypothesized causal pathway tested here, where *CHUK* (IKK-Alpha) serves as an instrumental variable that affects *NFKBIA* (IκB-Alpha), which in turn influences the expression of Gene X. (B) Schematic representing molecular mechanism of the NF-κB activation. (C) Instrumental Variable IV regression results, direction of *NFKBIA* effect (+/-), R-squared (R²), p-values (green highlighting indicates statistical significance), and weak instrument test results. (D) The diagram illustrates the comparison in Granger causality testing between two predictive models. The left panel shows the complete model where post-stimulation values of Gene X are predicted using both pre-stimulation *NFKBIA* and pre-stimulation Gene X values. The right panel represents the restricted model where post- stimulation Gene X values are predicted using only pre-stimulation Gene X values. The "vs" indicates the statistical comparison between these models; if the complete model significantly outperforms the restricted model, it suggests that *NFKBIA* Granger-causes Gene X, providing evidence for a temporal causal relationship. (E) The table illustrates statistical outcomes of Granger causality testing between *NFKBIA* and multiple inflammation-associated genes. The upper section displays results for "NFKBIA → X" (testing whether *NFKBIA* Granger-causes each gene), while the lower section shows "X → NFKBIA" (testing whether each tested gene Granger- causes *NFKBIA*). For each direction, both p-values and F-statistics are provided. Green- highlighted cells indicate statistically significant relationships (p < 0.05).

**Figure. S3.**
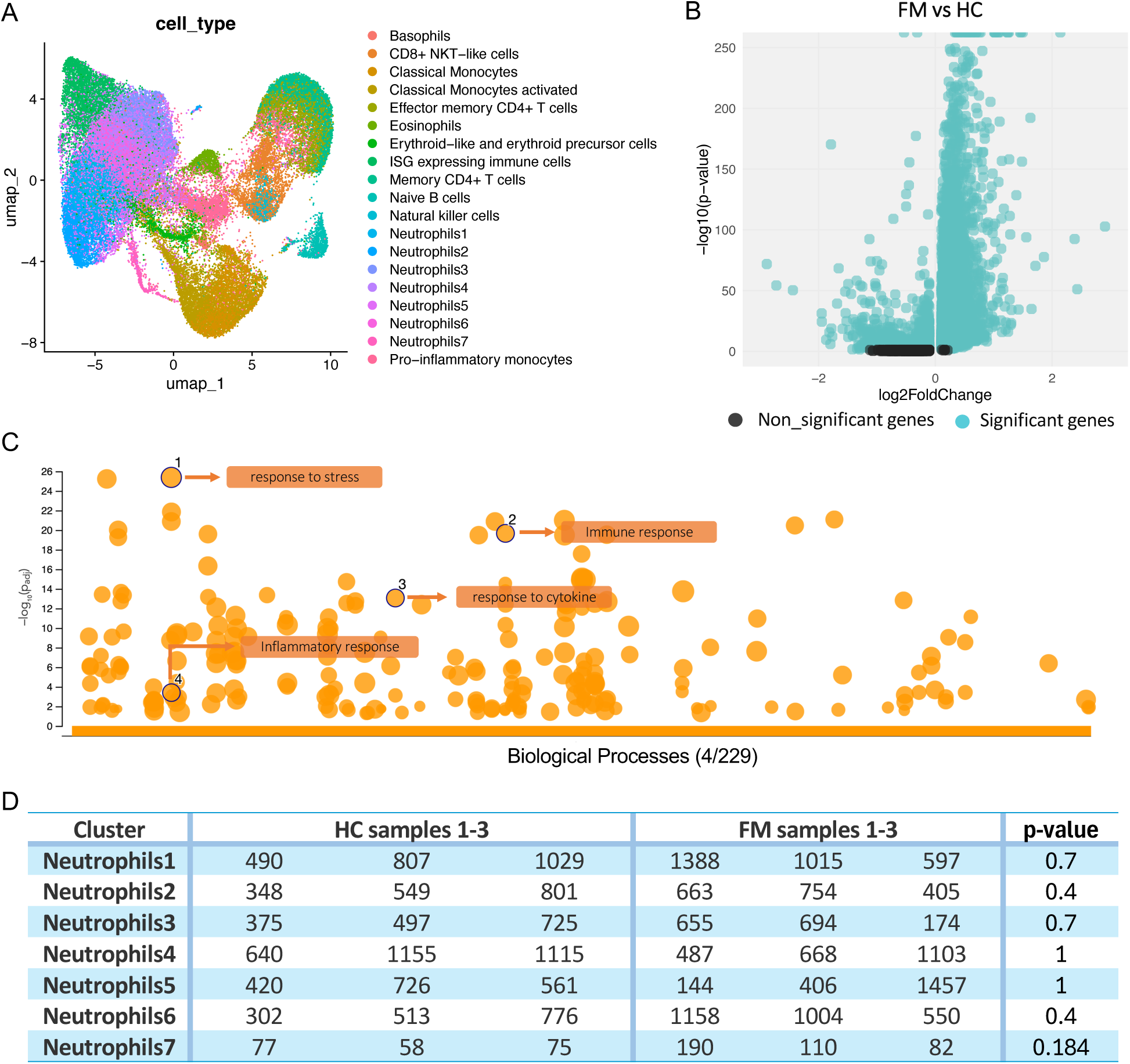
scRNA-seq identifies an elevated innate immune response pattern in the leukocytes of the whole blood of independent FM samples. (A) UMAP visualization depicting all the clusters identified through scRNA sequencing. (B) Scatterplot of the log2fold changes versus the statistical significance of the DEGs in neutrophils from FM patients versus HC. Black symbols represent genes regulated with an adjusted p-value of higher than 0.05, and blue symbols lower than p-value=0.05. (C) Statistics of Fgsea pathway analysis on DEGs in neutrophils from FM patients versus HC. (D) The number of cells in each neutrophil cluster in FM or HC samples.

**Figure. S4.**
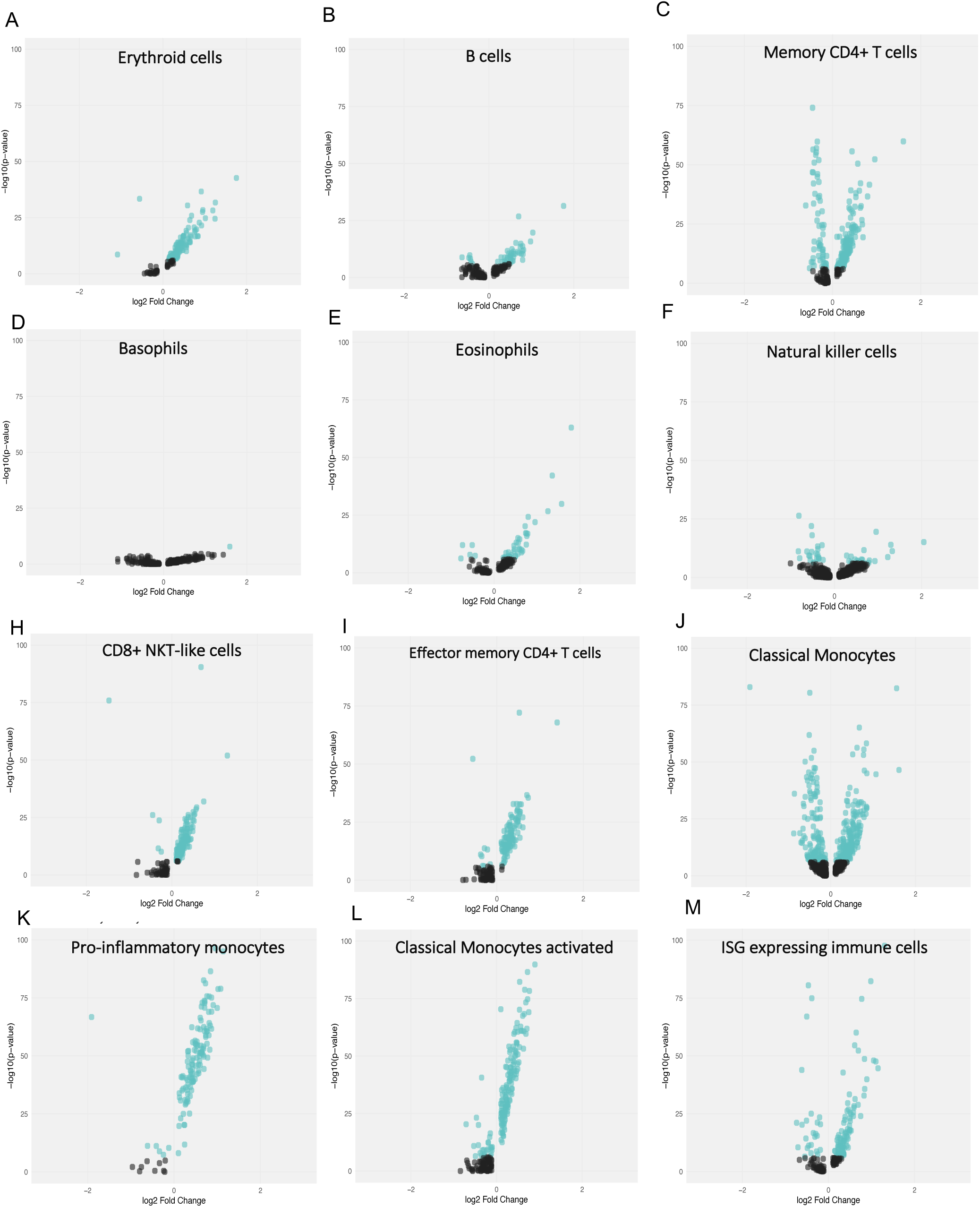
scRNA-seq identifies an up-regulated gene expression pattern in the leukocytes of whole blood of independent FM samples. (A-M) Scatterplot of the log2fold changes versus the statistical significance of the DEGs in each cell subtype when comparing FM versus HC.

**Figure. S5.**
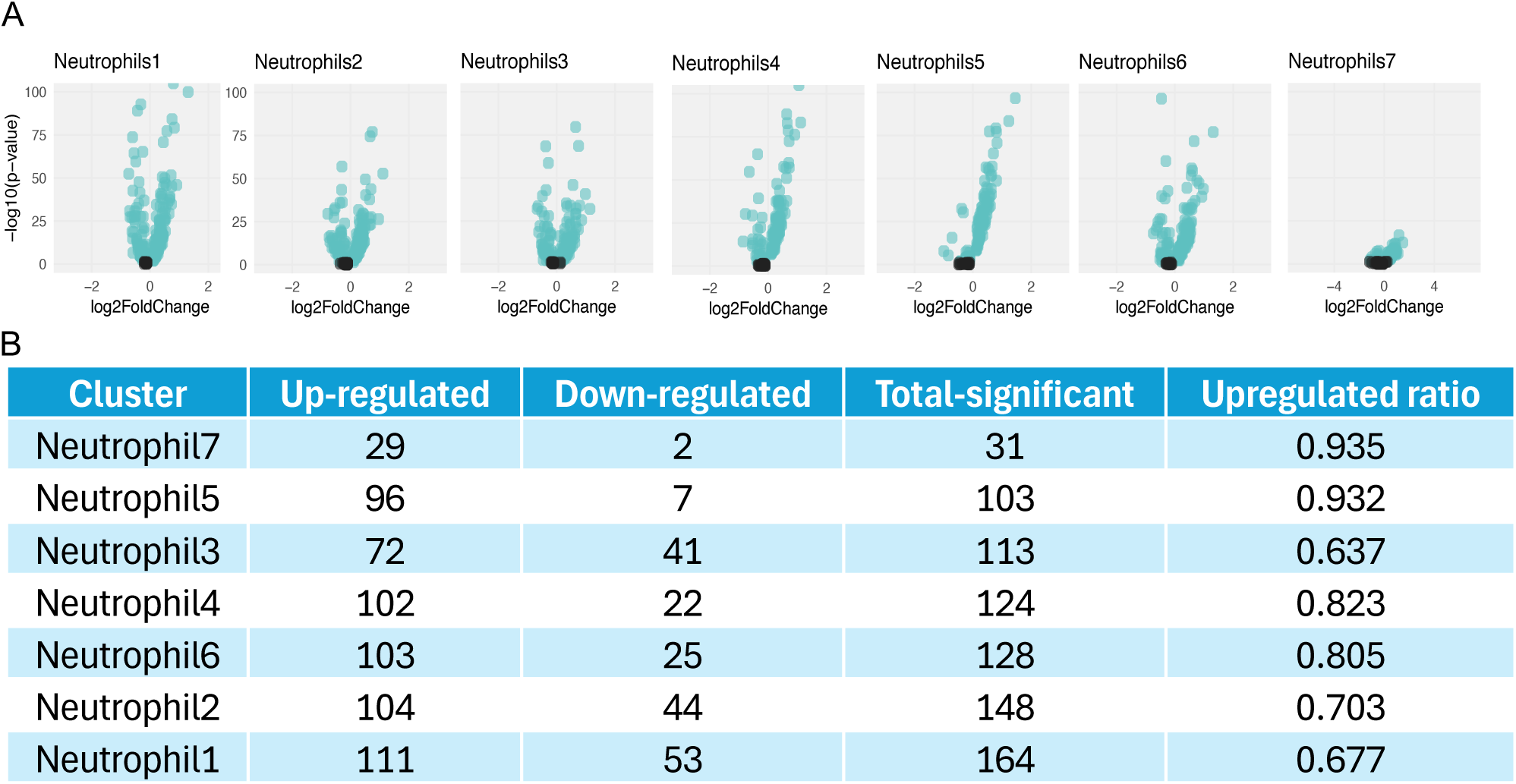
scRNA-seq identifies an up-regulated gene expression pattern in neutrophil subtypes in FM samples. (A) Scatterplot of the log2fold changes versus the statistical significance of the DEGs in each neutrophil subtype when comparing FM versus HC. (B) Table presenting the number of upregulated and downregulated genes, the total number of significantly differentially expressed genes, and the ratio of upregulated to total significantly differentially expressed genes when comparing FM patients versus HC in each neutrophil subcluster.

**Supplementary Table 1.** Differential gene expression analysis comparing non-stimulated neutrophils of FM patients versus HC. The "gene" column represents the names or identifiers of the genes analyzed. The "ENSID" column represents the corresponding Ensembl identifiers, which are unique references used in genomic databases. The "baseMean" column represents the average of the normalized counts for all samples, showing the mean expression level of each gene across the dataset. The "log2FoldChange" column represents the log2 fold change in expression between two conditions. The "lfcSE" column represents the standard error of the log2 fold change, estimating the variability of the log2 fold change. The "stat" column represents the test statistic used to determine the significance of the differential expression. The "pvalue" column represents the p-value for the test, indicating the probability that the observed difference in expression is due to chance. The "padj" column represents the p-value adjusted for multiple testing. The "Case" and "Control" columns represent the mean normalized counts for each group.

**Supplementary Table 2.** Fgsea pathway analysis results of DEG between non-stimulated neutrophils of FM patients versus HC. The "pathway" column represents the names or identifiers of the pathways analyzed, while the "desc" column represents a brief description of each pathway. The "pval" column represents the p-value for the enrichment test. The "padj" column represents the p-value adjusted for multiple testing. The "ES" column represents the enrichment score, showing the degree to which a gene set is overrepresented at the top or bottom of the ranked list of genes. The "NES" column represents the normalized enrichment score. The "nMoreExtreme" column represents the number of permutations where the observed enrichment score is as extreme or more extreme than the actual enrichment score observed. The "size" column represents the number of genes in the gene set. The "leadingEdge" column represents the genes that contribute most to the enrichment score.

**Supplementary Table 3.** gProfiler pathway analysis results of DEG between non-stimulated neutrophils of FM patients versus HC. The "term_name" column represents the name of the biological process term. The "term_id" column represents the unique identifier for the GO term. The "adjusted_p_value" column represents the p-value adjusted for multiple testing. The "negative_log10_of_adjusted_p_value" column represents the negative base-10 logarithm of the adjusted p-value. The "term_size" column represents the number of genes associated with the GO term in the entire database. The "query_size" column represents the number of genes from the input query that are associated with the GO term. The "effective_domain_size" column represents the number of genes in the background set that are associated with any GO term. The "intersections" column represents the specific genes from the query that intersect with the GO term.

**Supplementary Table 4.** Differential gene expression analysis comparing stimulated versus non-stimulated neutrophils from HC. The "gene" column represents the names or identifiers of the genes analyzed. The "ENSID" column represents the corresponding Ensembl identifiers, which are unique references used in genomic databases. The "baseMean" column represents the average of the normalized counts for all samples, representing the mean expression level of each gene across the dataset. The "log2FoldChange" column represents the log2 fold change in expression between two conditions. The "lfcSE" column represents the standard error of the log2 fold change, estimating the variability of the log2 fold change. The "stat" column represents the test statistic used to determine the significance of the differential expression. The "pvalue" column represents the p-value for the test, indicating the probability that the observed difference in expression is due to chance. The "padj" column represents the p-value adjusted for multiple testing. The "Stimulated" and "Non-stimulated" columns represent the mean normalized counts for each group.

**Supplementary Table 5.** Differential gene expression analysis comparing stimulated versus non-stimulated neutrophils from FM patients. The "gene" column represents the names or identifiers of the genes analyzed. The "ENSID" column represents the corresponding Ensembl identifiers, which are unique references used in genomic databases. The "baseMean" column represents the average of the normalized counts for all samples, representing the mean expression level of each gene across the dataset. The "log2FoldChange" column represents the log2 fold change in expression between two conditions. The "lfcSE" column represents the standard error of the log2 fold change, estimating the variability of the log2 fold change. The "stat" column represents the test statistic used to determine the significance of the differential expression. The "pvalue" column represents the p-value for the test, indicating the probability that the observed difference in expression is due to chance. The "padj" column represents the p-value adjusted for multiple testing. The "Stimulated" and "Non-stimulated" columns represent the mean normalized counts for each group.

**Supplementary Table 6.** Fgsea pathway analysis results of DEG between stimulated versus non-stimulated neutrophils from HC. The "pathway" column represents the names or identifiers of the pathways analyzed, while the "desc" column represents a brief description of each pathway. The "pval" column represents the p-value for the enrichment test. The "padj" column represents the p-value adjusted for multiple testing. The "ES" column represents the enrichment score, representing the degree to which a gene set is overrepresented at the top or bottom of the ranked list of genes. The "NES" column represents the normalized enrichment score. The "nMoreExtreme" column represents the number of permutations where the observed enrichment score is as extreme or more extreme than the actual enrichment score observed. The "size" column represents the number of genes in the gene set. The "leadingEdge" column represents the genes that contribute most to the enrichment score.

**Supplementary Table 7.** Fgsea pathway analysis results of DEG between stimulated versus non-stimulated neutrophils from FM patients. The "pathway" column represents the names or identifiers of the pathways analyzed, while the "desc" column represents a brief description of each pathway. The "pval" column represents the p-value for the enrichment test. The "padj" column represents the p-value adjusted for multiple testing. The "ES" column represents the enrichment score, representing the degree to which a gene set is overrepresented at the top or bottom of the ranked list of genes. The "NES" column represents the normalized enrichment score. The "nMoreExtreme" column represents the number of permutations where the observed enrichment score is as extreme or more extreme than the actual enrichment score observed. The "size" column represents the number of genes in the gene set. The "leadingEdge" column represents the genes that contribute most to the enrichment score.

**Supplementary Table 8.** gProfiler pathway analysis results of DEG between stimulated versus non-stimulated neutrophils from HC. The "term_name" column represents the name of the biological process term. The "term_id" column represents the unique identifier for the GO term. The "adjusted_p_value" column represents the p-value adjusted for multiple testing. The "negative_log10_of_adjusted_p_value" column represents the negative base-10 logarithm of the adjusted p-value. The "term_size" column represents the number of genes associated with the GO term in the entire database. The "query_size" column represents the number of genes from the input query that are associated with the GO term. The "effective_domain_size" column represents the number of genes in the background set that are associated with any GO term. The "intersections" column represents the specific genes from the query that intersect with the GO term.

**Supplementary Table 9.** gProfiler pathway analysis results of DEG between stimulated versus non-stimulated neutrophils from FM patients. The "term_name" column represents the name of the biological process term. The "term_id" column represents the unique identifier for the GO term. The "adjusted_p_value" column represents the p-value adjusted for multiple testing. The "negative_log10_of_adjusted_p_value" column represents the negative base-10 logarithm of the adjusted p-value. The "term_size" column represents the number of genes associated with the GO term in the entire database. The "query_size" column represents the number of genes from the input query that are associated with the GO term. The "effective_domain_size" column represents the number of genes in the background set that are associated with any GO term.

**Supplementary Table 10.** Interaction analysis on FM patients/HC/stimulated/non- stimulated conditions. The "gene" column represents the names or identifiers of the genes analyzed. The "ENSID" column represents the corresponding Ensembl identifiers, which are unique references used in genomic databases. The "baseMean" column represents the average of the normalized counts for all samples, representing the mean expression level of each gene across the dataset. The "log2FoldChange" column represents the log2 fold change in expression between two conditions. The "lfcSE" column represents the standard error of the log2 fold change, estimating the variability of the log2 fold change. The "stat" column represents the test statistic used to determine the significance of the differential expression. The "pvalue" column represents the p-value for the test, indicating the probability that the observed difference in expression is due to chance. The "padj" column represents the p-value adjusted for multiple testing. The last four columns represent the mean normalized counts for each group.

**Supplementary Table 11.** Differential gene expression analysis comparing stimulated versus non-stimulated neutrophils from FM Improvers. The "gene" column represents the names or identifiers of the genes analyzed. The "ENSID" column represents the corresponding Ensembl identifiers, which are unique references used in genomic databases. The "baseMean" column represents the average of the normalized counts for all samples, representing the mean expression level of each gene across the dataset. The "log2FoldChange" column represents the log2 fold change in expression between two conditions. The "lfcSE" column represents the standard error of the log2 fold change, estimating the variability of the log2 fold change. The "stat" column represents the test statistic used to determine the significance of the differential expression. The "pvalue" column represents the p-value for the test, indicating the probability that the observed difference in expression is due to chance. The "padj" column represents the p-value adjusted for multiple testing. The "Stimulated" and "Non-stimulated" columns represent the mean normalized counts for each group.

**Supplementary Table 12.** Differential gene expression analysis comparing stimulated versus non-stimulated neutrophils from FM Persisters. The "gene" column represents the names or identifiers of the genes analyzed. The "ENSID" column represents the corresponding Ensembl identifiers, which are unique references used in genomic databases. The "baseMean" column represents the average of the normalized counts for all samples, representing the mean expression level of each gene across the dataset. The "log2FoldChange" column represents the log2 fold change in expression between two conditions. The "lfcSE" column represents the standard error of the log2 fold change, estimating the variability of the log2 fold change. The "stat" column represents the test statistic used to determine the significance of the differential expression. The "pvalue" column represents the p-value for the test, indicating the probability that the observed difference in expression is due to chance. The "padj" column represents the p-value adjusted for multiple testing. The "Stimulated" and "Non-stimulated" columns represent the mean normalized counts for each group.

**Supplementary Table 13.** Summary of medication category usage among Improvers and Persisters. The table represents the number of individuals in each group (Improvers vs. Persisters) who reported using drugs within each category.

**Supplementary Table 14.** Association of FM duration with symptom improvement. The distribution of FM duration (in years) among Improvers and Persisters, along with the results of Welch’s t-test and Wilcoxon rank-sum test evaluating group differences.

**Supplementary Table 15.** Transcription factor analysis on the results of differential gene expression in non-stimulated neutrophils of FM patients versus HC. The "term_name" column represents the name of the transcription factor and its associated motif. The "term_id" column represents the unique identifier for the transcription factor term. The "adjusted_p_value" column represents the p-value adjusted for multiple testing. The "negative_log10_of_adjusted_p_value" column represents the negative base-10 logarithm of the adjusted p-value. The "term_size" column represents the number of genes associated with the transcription factor in the entire database. The "query_size" column represents the number of genes from the input query that are associated with the transcription factor. The "intersection_size" column represents the number of genes from the query that intersect with the transcription factor term. The "effective_domain_size" column represents the number of genes in the background set that are associated with any transcription factor term. The "intersections" column represents the specific genes from the query that intersect with the transcription factor term. Only significant results are presented.

**Supplementary Table 16.** Transcription factor analysis on interaction term analysis results on FM patients/HC/stimulated/non-stimulated conditions. The "term_name" column represents the name of the transcription factor and its associated motif. The "term_id" column represents the unique identifier for the transcription factor term. The "adjusted_p_value" column represents the p-value adjusted for multiple testing. The "negative_log10_of_adjusted_p_value" column represents the negative base-10 logarithm of the adjusted p-value. The "term_size" column represents the number of genes associated with the transcription factor in the entire database. The "query_size" column represents the number of genes from the input query that are associated with the transcription factor. The "intersection_size" column represents the number of genes from the query that intersect with the transcription factor term. The "effective_domain_size" column represents the number of genes in the background set that are associated with any transcription factor term. The "intersections" column represents the specific genes from the query that intersect with the transcription factor term. Only significant results are presented.

**Supplementary Table 17.** Differential gene expression analysis comparing scRNA seq data of FM patients versus HC regardless of cell type. The "gene" column represents the analyzed genes. The "p_val" column represents the raw p-value for differential expression significance. The "avg_log2FC" column represents the average log2 fold change in expression between FM patients and HC, where a positive value signifies higher expression in FM patients. The "pct.1" and "pct.2" columns represent the proportion of cells expressing each gene in FM patients and HC, respectively. The "p_val_adj" column represents the p-value adjusted for multiple testing.

**Supplementary Table 18.** gProfiler pathway analysis results of DEG between FM patients and HC in sc RNA seq data, regardless of cell type. The "term_name" column represents the name of the biological process term. The "term_id" column represents the unique identifier for the term. The "adjusted_p_value" column represents the p-value adjusted for multiple testing. The "negative_log10_of_adjusted_p_value" column represents the negative base-10 logarithm of the adjusted p-value. The "term_size" column represents the number of genes associated with the term in the entire database. The "query_size" column represents the number of genes from the input query that are associated with the term. The "intersection_size" column represents the number of overlapping genes between the input query and the term. The "effective_domain_size" column represents the number of genes in the background set associated with any term. The "intersections" column represents the specific genes from the input query that intersect with the term.

**Supplementary Table 19.** Differential gene expression analysis comparing sc RNA seq data of FM patients versus HC in neutrophils regardless of neutrophil subtype. The "gene" column represents the analyzed genes. The "p_val" column represents the raw p-value for differential expression significance. The "avg_log2FC" column represents the average log2 fold change in expression between FM patients and HC, where a positive value signifies higher expression in FM patients. The "pct.1" and "pct.2" columns represent the proportion of cells expressing each gene in FM patients and HC, respectively. The "p_val_adj" column represents the p-value adjusted for multiple testing.

**Supplementary Table 20.** gProfiler pathway analysis results for differentially DEG between FM patients and HC in sc RNA seq data in neutrophils regardless of neutrophil subtype. The "term_name" column represents the name of the biological process term. The "term_id" column represents the unique identifier for the term. The "adjusted_p_value" column represents the p-value adjusted for multiple testing. The "negative_log10_of_adjusted_p_value" column represents the negative base-10 logarithm of the adjusted p-value. The "term_size" column represents the number of genes associated with the term in the entire database. The "query_size" column represents the number of genes from the input query that are associated with the term. The "intersection_size" column represents the number of overlapping genes between the input query and the term. The "effective_domain_size" column represents the number of genes in the background set associated with any term. The "intersections" column represents the specific genes from the input query that intersect with the term.

**Supplementary Table 21.** Differential gene expression analysis comparing sc RNA seq data of FM patients versus HC in each neutrophil subtype. The "gene" column represents the analyzed genes. The "p_val" column represents the raw p-value for differential expression significance. The "avg_log2FC" column represents the average log2 fold change in expression between FM patients and HC, where a positive value signifies higher expression in the cluster mentioned in the "cluster" column. The "pct.1" and "pct.2" columns represent the proportion of cells expressing each gene in FM patients and HC, respectively. The "p_val_adj" column represents the p-value adjusted for multiple testing.

**Supplementary Table 22.** gProfiler pathway analysis results of DEG between FM patients and HC in scRNA seq data, for each cell type. The "term_name" column represents the name of the biological process term. The "term_id" column represents the unique identifier for the term. The "adjusted_p_value" column represents the p-value adjusted for multiple testing for each cell type/subtype separately.

**Supplementary Table 23.** Differential gene expression analysis comparing each neutrophil subtype versus all other neutrophil subtypes in scRNA seq data. **The "gene" column represents** the analyzed genes. The "p_val" column represents the raw p-value for differential expression significance. The "avg_log2FC" column represents the average log2 fold change in expression of the cluster, where a positive value signifies higher expression of the cluster mentioned in the "cluster" column. The "pct.1" and "pct.2" columns represent the proportion of cells expressing each gene in each subtype versus others. The "p_val_adj" column represents the p-value adjusted for multiple testing.

**Supplementary Table 24.** A collection of neutrophil markers that were deemed significant in each neutrophil subtype in scRNA-seq data. The "Cluster" column represents the neutrophil subtype. The "Marker" column represents both the gene name and the conventional name for each marker. "p_val_adj" represents the adjusted p-value for significance as a cluster marker. "avg_log2fc" represents the log2 fold change. The "Direction as cluster marker" column represents whether the marker is upregulated or downregulated within the respective cluster. "Direction in FM" represents how the gene is regulated when comparing that cluster in FM versus HC. "Adjusted p-value for FM vs HC" represents the statistical significance of this comparison. The "Description" column represents additional details about each marker.

